# Distinct SARS-CoV-2 sensing pathways in pDCs driving TLR7-antiviral vs. TLR2-immunopathological responses in COVID-19

**DOI:** 10.1101/2021.11.23.469755

**Authors:** Renée M. van der Sluis, Lamin B. Cham, Albert Gris Oliver, Kristine R. Gammelgaard, Jesper G. Pedersen, Manja Idorn, Ulvi Ahmadov, Sabine Sanches Hernandez, Ena Cémalovic, Stine H. Godsk, Jacob Thyrsted, Jesper D. Gunst, Silke D. Nielsen, Janni J. Jørgensen, Tobias Wang Bjerg, Anders Laustsen, Line S. Reinert, David Olagnier, Rasmus O. Bak, Mads Kjolby, Christian K. Holm, Martin Tolstrup, Søren R. Paludan, Lasse S. Kristensen, Ole S. Søgaard, Martin R. Jakobsen

## Abstract

Understanding the molecular pathways driving the acute antiviral and inflammatory response to SARS-CoV-2 infection is critical for developing treatments for severe COVID-19. Here we show that in COVID-19 patients, circulating plasmacytoid dendritic cells (pDCs) decline early after symptom onset and this correlated with COVID-19 disease severity. This transient depletion coincides with decreased expression of antiviral type I IFNα and the systemic inflammatory cytokines CXCL10 and IL-6. Importantly, COVID-19 disease severity correlated with decreased pDC frequency in peripheral blood. Using an *in vitro* stem cell-based human pDC model, we demonstrate that pDCs directly sense SARS-CoV-2 and in response produce multiple antiviral (IFNα and IFNλ1) and inflammatory (IL-6, IL-8, CXCL10) cytokines. This immune response is sufficient to protect epithelial cells from de novo SARS-CoV-2 infection. Targeted deletion of specific sensing pathways identified TLR7-MyD88 signaling as being crucial for production of the antiviral IFNs, whereas TLR2 is responsible for the inflammatory IL-6 response. Surprisingly, we found that SARS-CoV-2 engages the neuropilin-1 receptor on pDCs to mitigate the antiviral IFNs but not the IL-6 response. These results demonstrate distinct sensing pathways used by pDCs to elicit antiviral vs. immunopathological responses to SARS-CoV-2 and suggest that targeting neuropilin-1 on pDCs may be clinically relevant for mounting TLR7-mediated antiviral protection.

**One Sentence Summary:** pDCs sense SARS-CoV-2 and elicit antiviral protection of lung epithelial cells through TLR7, while recognition of TLR2 elicits an IL-6 inflammatory response associated with immunopathology.

**Graphical abstract:**
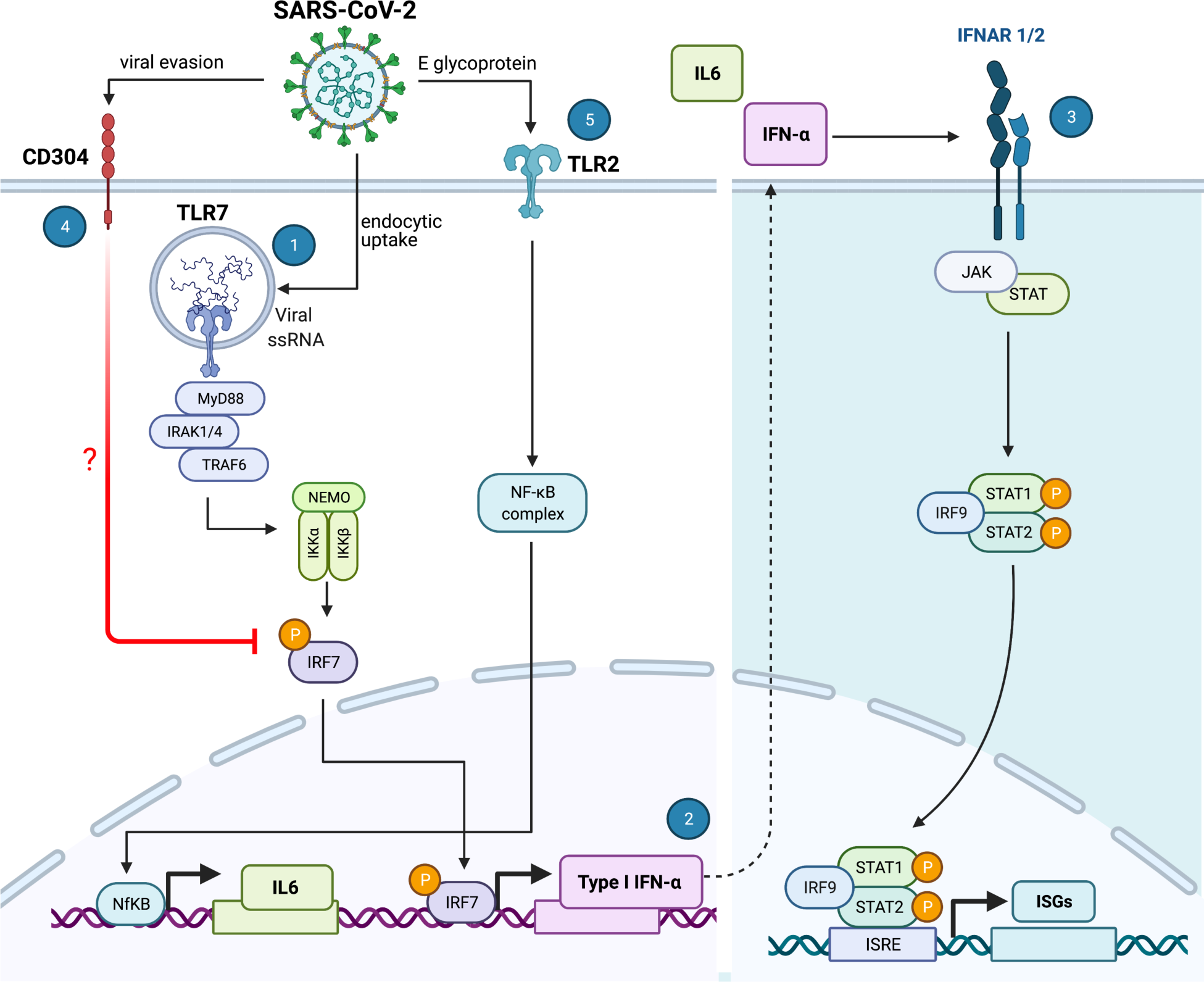
SARS-CoV-2 sensing by plasmacytoid dendritic cells. SARS-CoV-2 is internalized by pDCs via a yet unknown endocytic mechanism. The intracellular TLR7 sensor detects viral RNA and induces a signaling cascade involving MyD88-IRAK4-TRAF6 (1) to induce CXCL10 and, via IRF7 phosphorylation and translocation, inducing type I and III Interferons (2). Once secreted, type I and III IFNs initiate autocrine and paracrine signals that induce the expression of IFN stimulated genes (ISGs), thereby facilitating an antiviral response that can protect the cell against infection. However, SARS-CoV-2, has the intrinsic property to facilitate CD304 signaling, potentially by interfering with IRF7 nuclear translocation, thereby inhibiting type I IFNα production and thus reducing the antiviral response generated by the pDC (4). Furthermore, the SARS-CoV-2 envelope (E) glycoprotein is sensed by the extracellular TLR2/6 heterodimer and this facilitates production of the inflammatory IL-6 cytokine (5). Illustration was created with BioRender.com

## Introduction

The severe acute respiratory syndrome coronavirus-2 (SARS-CoV-2) has, since its first appearance in 2019, resulted in a devastating pandemic of coronavirus disease 2019 (COVID-19) that prevails mid 2021 (*1, 2*). The severity of COVID-19 is highly variable between individuals and a great effort is made to understand why some people develop mild disease whilst others require hospitalization (*3, 4*). A reported driver of disease severity is the imbalanced induction of an immune response consisting of a broad range of inflammatory cytokines combined with a delayed induction of antiviral interferons (IFNs) (*5-7*). Factors associated with severe disease are inborn errors in the Toll-like receptor (TLR)3 and interferon regulatory factor (IRF)7-dependent type I IFN production and the presence of auto-antibodies against type I IFNs (*8, 9*). This indicates that sufficient amounts of IFNs are essential for controlling the infection. Yet, it remains unclear which immune cells detect SARS-CoV-2 and initiate the inflammatory response. Alveolar macrophages seem incapable of sensing SARS-CoV-2 (*10*) and whether in vitro generated macrophages and myeloid dendritic cells (DCs) are able to elicit production of pro-inflammatory and antiviral cytokines in response to SARS-CoV-2 is currently unclear (*11, 12*); and one study suggests that lung epithelial cells are needed for the macrophages to produce antiviral cytokines (*13*). Importantly, lung epithelial cells can however detect SARS-CoV2 and produce type I IFNβ and type III IFNλ1, but only after initiation of virus replication (*14, 15*).

Plasmacytoid DCs (pDCs) are an autonomous cell type and major producers of type I IFNα, making them pivotal for the human immune system to control viral infections (*16*). Clinical studies reveal that severe COVID-19 cases have a reduction in circulating pDCs as well as minimal influx of pDCs into the lungs compared to patients with moderate disease and healthy controls (*6, 17-22*). These severe cases of COVID-19 also exhibited reduced type I IFNα, type III IFNλ and interleukin (IL-)3 levels in plasma, of which IL-3 is known to be important for pDC function (*19*). Whether disease severity is due to the lack of pDCs in the lungs or due to dysfunctional cytokine production by the pDCs, remains unclear. Furthermore, the mechanism of how pDCs may sense SARS-CoV-2 has not been resolved.

Generally, cytokine production from pDCs is triggered upon the innate detection of viral components via various extra- and intra-cellular receptors also known as pattern recognition receptors (PRRs). In particular, the Toll-like receptors (TLRs) and retinoic acid-inducible gene I (RIG-I)-like receptors (RLR) are the major receptor classes responsible for sensing RNA virus infection and triggering antiviral IFN production (*23*). In the present study, we explored via which molecular mechanism human pDCs sense SARS-CoV-2, by using a CRISPR-editing approach to screen for several innate immune sensor pathways that are required for the production of antiviral IFNs and inflammatory cytokines upon viral sensing.

## RESULTS

### Circulating pDCs numbers are linked to systemic inflammatory signals during SARS-CoV-2 infection

To investigate the impact of SARS-CoV-2 infection on frequency and phenotype of circulating pDCs, we collected blood samples from patients hospitalized for COVID-19 at hospital admission (day 1) and 5 days after admission. We categorized patients according to symptom duration, which was defined as time from onset of the first self-reported COVID-19 symptom to date of hospital admission (0 to 4; 5 to 8; 9 to 12; and ≥13 days). When we compared the percentage of pDCs out of total PBMCs across the different symptom duration categories, we found a significant lower frequency of circulating pDCs among patients with symptom duration between 5 to 12 days as compared to those with short symptom duration (0 to 4 days) and long symptom duration (≥13 days) (Figure 1A, EV1A), which was not observed for the myeloid DC subset (Figure 1B). We next explored the changes of pDC percentage over time within each patient and observed a significant decrease in pDC frequency and numbers after 5 days of hospitalization (Figure 1C, EV1B), which was not seen in the myeloid DC subset (Figure 1D). Compared to healthy controls (HC), pDC frequency and counts in peripheral blood were decreased in COVID-19 patients (Figure EV1C-D). We then evaluated whether the reduction in circulating pDCs was associated with systemic inflammatory cytokine levels. Except for IL-8, the plasma concentration of multiple pro-inflammatory cytokines, in particularly IFNα2a, CXCL10 and IL-6, changed significantly between D1 and D5 (Figure 1E-J). To determine the association between pDC frequency (in total PBMC) and disease severity, we next categorized patients into i) Hospitalized and no oxygen supplementation required, ii) Hospitalized and nasal oxygen supplementation required, and iii) Hospitalized and high flow oxygen supplementation required. The pDC frequency was significantly lower among patients who required high flow oxygen supplementation as compared to the group that did not require oxygen supplementation (Figure 1K) and a correlation between decreased pDC frequency and disease severity was observed (Figure 1L). These findings suggest that COVID-19 disease can be associated with a decrease in the percentage of pDCs in peripheral blood and that pDCs are a driver of the inflammatory signals observed during acute SARS-CoV-2 infection.

**Figure 1.**
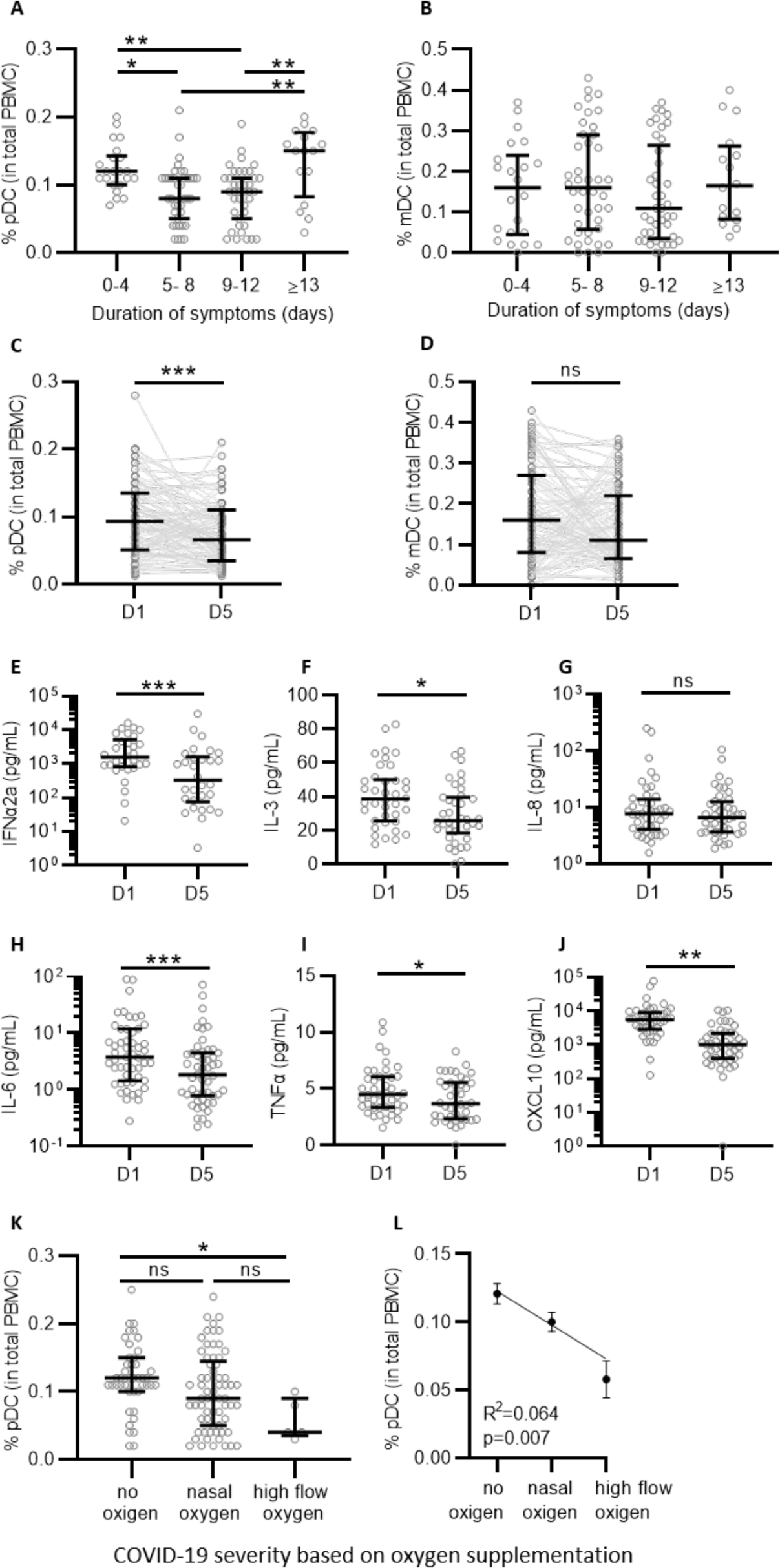
Infection with SARS-CoV-2 is associated with a decrease in plasmacytoid dendritic cells. PBMCs and plasma samples were collected from SARS-CoV-2 infected individuals at time of hospitalization (D1) and after 5 days (D5). Percentage of pDCs (A) and mDCs (B) from total PBMCs collected at D1 was quantified by flow and grouped to self-reported duration of symptoms prior to D1 (A; 0-4 n= 22; 5-8 n=39; 9-12 n= 39; ≥13 n=16). Percentage of pDCs (C) and mDCs (D) at D1 and D5 matched samples from all patients independent of symptom duration (n=93). E-J: Matched D1 and D5 plasma samples were available from 55 donors and protein levels were determined for IFNα2a (E, n=30), IL-3 (F, n=38), IL-8 (G, n=44), IL-6 (H, n=51), TNFα (I, n=42) and CXCL10 (J, n=46), note that some samples could not be quantified and were excluded from the analysis. Percentage of pDCs grouped by COVID-19 severity, as determined by oxygen supplementation (K, no oxygen n=43, nasal oxygen n=65 and high flow oxygen n=5). Percentage of pDCs was correlated to COVID-19 disease severity (L) using simple linear regression. Each dot represents a patient, lines with error bars show the median values with interquartile ranges. Statistical significance was determined using the Kruskal-Wallis test (A-B), Wilcoxon matched pairs signed rank test (B-K) and simple linear regression (L). *<p0.05, **<p0.01 ***<p0.001, ns= not significant.

### Human pDCs sense SARS-CoV-2 but are refractory to infection

Studying viral sensing by human pDCs is hampered by the limited amount of pDCs that can be obtained from peripheral blood and their incapability to be genetically modified. To overcome this and enable the investigation of potential pDC sensing mechanisms of SARS-CoV-2, we adopted a cellular platform designed to generate human primary pDCs *ex vivo* using hematopoietic stem and progenitor cells (HSPC) from healthy individuals (Appendix Figure S1) (*24, 25*). The HSPC-derived pDCs, produced from multiple healthy donors, were exposed to two different SARS-CoV-2 isolates at 0.1 multiplicity of infection (MOI, as determined by limiting dilution on VeroE6-TMPRSS2 cells); the Freiburg isolate (FR2020) which is an early 2020 Wuhan-like strain and the SARS-CoV-2 alpha variant (B.1.1.7). Type I IFNα and CXCL10 production was assessed longitudinally and found to be induced by both variants (Figure 2A-B) with a trend towards a more rapid type I IFNα induction observed for the SARS-CoV-2 alpha variant. To further characterize how pDCs sense SARS-CoV-2, we continued with the SARS-CoV-2 Freiburg isolate. First, pDCs were exposed to TLR agonists or SARS-CoV-2 and after 24hrs cell culture supernatants and pDCs were collected to assess a broader range of inflammatory cytokines, at both the protein and mRNA levels respectively. Following SARS-CoV-2 exposure, we observed increased production of type I IFNα and type III IFNλ1, but not type I IFNβ and type II IFNγ (Figure 2C-F), resembling the TLR7 agonist response. The cytokines IL-6, IL-8 and CXCL10 were likewise induced, with SARS-CoV-2 inducing higher IL-6 levels than the TLR7 agonist (Figure 2G-I). Tumor necrosis factor (TNF)α was only marginally induced in pDCs from some donors challenged with SARS-CoV-2 (Figure 2J), a pattern resembling neither TLR3 nor TLR7 agonists. Next, we evaluated the type I IFNα expression pattern relative to viral titer and duration of exposure. A viral MOI of 1, resulted in a strong type I IFNα response on both RNA and protein levels (Figure 2K-L), and a clear positive correlation between type I IFNα induction and exposure time was observed (Figure 2M). A similar pattern was observed for type III IFNλ1 and multiple inflammatory cytokines (Figure EV2). The HSPC-pDC cytokine responses to SARS-CoV-2 were very similar to what was obtained when using freshly isolated human pDCs from peripheral blood (Figure EV3). Next, we assessed whether SARS-CoV-2 was able to replicate in pDCs. However, no viral products indicative of SARS-CoV-2 replication were detected in pDCs (Appendix Figure S2), which is supported by others (*26*). Overall, these results demonstrate that pDCs are capable of sensing SARS-CoV-2 and in response produce type I IFNα and numerous inflammatory cytokines that are important to the cytokine storm observed in people suffering from severe COVID-19 disease (*5-7*).

**Figure 2.**
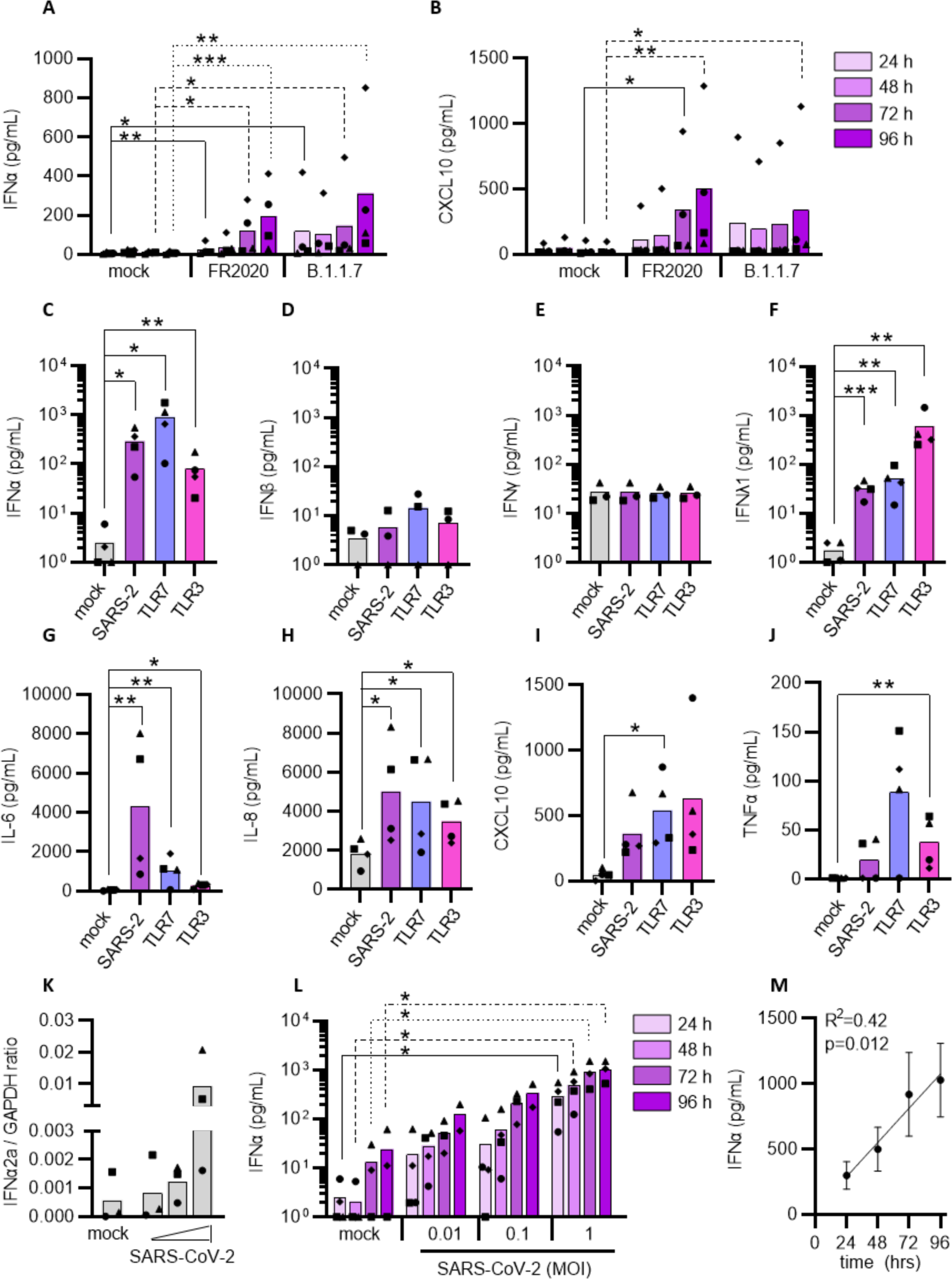
Plasmacytoid DCs can sense SARS-CoV-2 and induce an inflammatory response. pDCs were either mock treated or exposed to the SARS-CoV-2 FR2020 early Wuhan-like strain or the SARS-CoV-2 alpha variant B.1.1.7 (0.1 MOI). Supernatants were collected at indicated time points and the production of type I IFNα (A) and CXCL10 (B) was quantified. The FR2020 strain was used in subsequent experiments where pDC were either mock treated (mock, grey), exposed to SARS-CoV-2 at 1 MOI (SARS-2, purple), TLR7 (2.5 μg/mL R837, blue) or TLR3 agonist (800 ng/mL poly(I:C), pink). Supernatants were collected after 24 hrs and analyzed for type I IFNα (C), IFNβ (D), type II IFNγ (E), type III IFNλ1 (F), IL-6 (G), IL-8 (H), CXCL10 (I) and TNFα (J) expression by ELISA. To evaluate the cytokine response to viral titers and exposure duration, pDCs were exposed to increasing viral inoculums (MOI of 0.01, 0.1 and 1) and IFNα2a mRNA expression was quantified at 24 hrs (K) and IFNα protein secretion at 24, 48, 72 and 96 hrs (L). Graph depicting simple linear regression of IFNα protein with time of exposure (M). Bars and lines represent mean values and symbols represent individual pDC donors (n=3-4). Equal symbols represent equal donors (A-B and C-L). Statistical significance was determined using the ratio paired student T test and compared the treated condition with the time point-matched mock condition (A-L) and simple linear regression (M). *<p0.05, **<p0.01 ***<p0.001.

### Detection of SARS-CoV-2 in pDCs facilitates a protective antiviral response through a broad inflammatory gene signature

A hallmark of antiviral activity is protection of target cells against the pathogen. To investigate if pDC-secreted cytokines protected cells from SARS-CoV-2 infection, we next exploited two different lung epithelial cell types - A549 hACE2 and Calu-3 – and exposed them to cell culture supernatant from pDCs that were either cultured as normal or exposed to SARS-CoV-2, followed by virus inoculation. Pre-treatment with supernatant from SARS-CoV-2-exposed pDCs reduced virus replication in both cell lines in a dose dependent manner (Figure EV4A-B). Blocking type I IFN signaling enhanced virus replication for all pDC donors tested, but this did not reach statistical significance, indicating that protection was mediated partially by type I IFNs (Figure EV4C). Overall, these data indicate that cytokines produced by pDCs in response to SARS-CoV-2 can protect lung cells from infection by reducing virus replication and thereby limit viral spread.

To broader investigate the nature and timing of SARS-CoV-2-induced antiviral responses in pDCs we next profiled 789 selected genes covering major immunological pathways (Table S1) using the NanoString nCounter technology (*27*). We profiled the selected genes 4, 24 and 48 hrs after SARS-CoV-2 infection in two individual donors found to be high (D^high^) and low (D^low^) responders in terms of type I IFNα production (see Figure 2L, where triangles denote D^high^ and squares denote D^low^). There was a large overlap between the two donors in gene expression detected above background levels (Figure EV5A) and while multiple genes were induced as early as 4 hrs post SARS-CoV-2 exposure, the immunological response seemed stronger after 48 hrs (Figure 3A-B, Figure EV5B-D, Table S2, Table S3). Interestingly, IL-6, CXCL10, CCL2 and CCL8 were among the most upregulated genes in both donors after both 4 and 48 hrs (Figure EV5E-F). When comparing the expression of the most strongly induced genes (fold change > 2, relative to mock treated cells) after 48 hours of infection (104 genes in D^high^ and 66 genes in D^low^) with the earlier time points, it became apparent that distinct gene sets behaved differently (Figure 3C-D). For instance, some genes peaked at 4 hrs or 24 hrs, some clearly peaked at 48 hrs (cluster 4 Figure 3C and cluster 5 Figure 3D), while others had a clear biphasic expression format (induced early, disappearing after 24 hrs and then re-induced after 48 hrs; cluster 6 Figure 3C and cluster 2 Figure 3D). These gene clusters included pathways involved in the pDCs’ anti-viral response, and pathways representing more general innate immune activation (Appendix Figure S3 and S4). To confirm the intriguing biphasic gene induction, CXCL10 gene expression was selected for further analysis in multiple pDC donors. RT-qPCR analysis revealed that the wave pattern of gene induction was specific for SARS-CoV-2 sensing and did not follow a TLR7 or TLR3 agonist-induced pattern (Figure 3E), confirming our previous observations. Although the NanoString analysis would ideally have been performed using more donors, the data do demonstrate that SARS-CoV-2 activates different steps at different time points in the pDCs’ viral sensory pathways, indicating a multifaceted sensory mechanism, where antiviral type I IFNα is found in the early phase, succeeded by excessive inflammatory cytokines at later times. This reflects in part the pathogenesis of COVID-19 (*7*).

**Figure 3.**
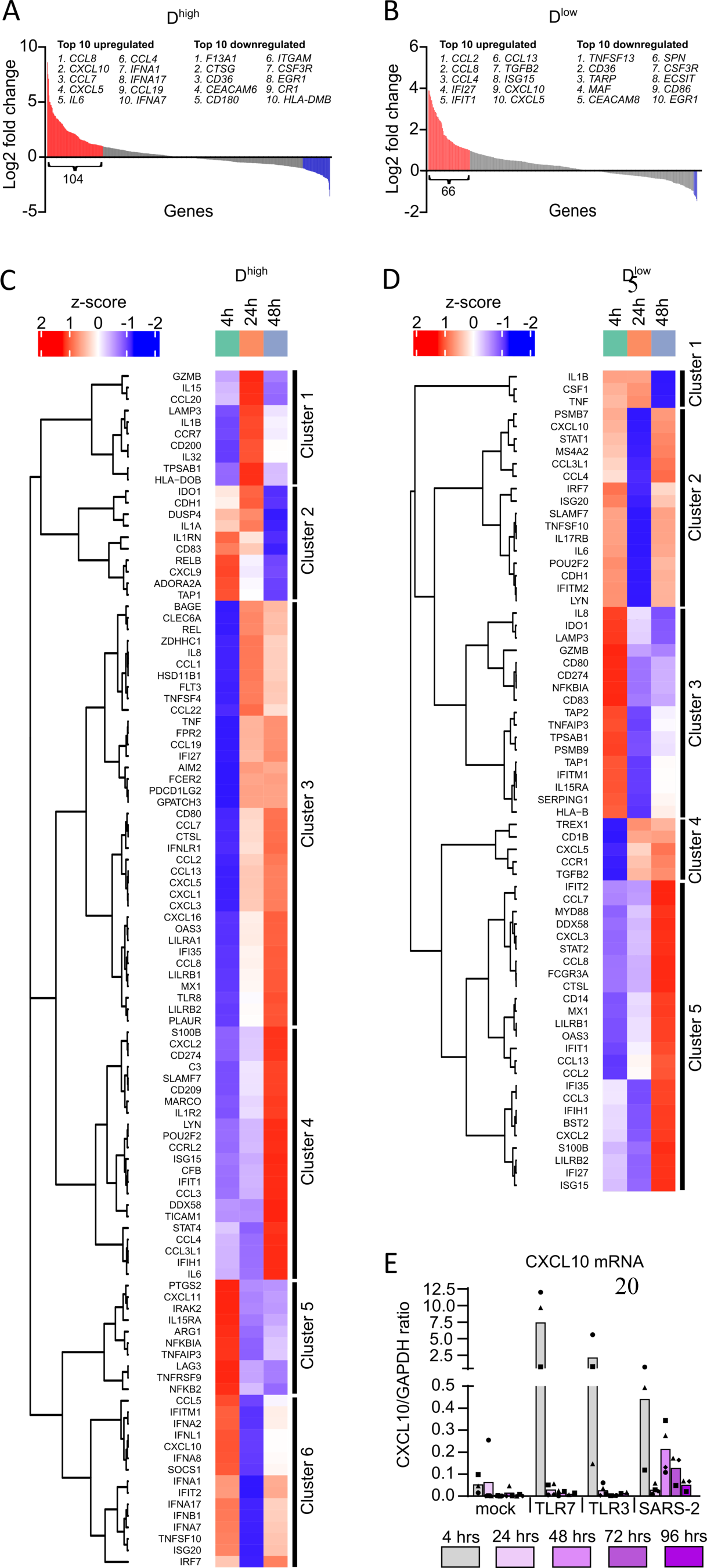
Nature and timing of SARS-CoV-2-induced gene expression changes in pDCs. Waterfall plots illustrating gene expression changes in pDCs 48 hrs post SARS-CoV-2 exposure, relative to mock treated cells, from two donors; D^high^ (A) and D^low^ (B), indicating genes with >2 fold change in red (upregulated) and blue (downregulated). Heat maps and unsupervised hierarchical cluster analyses of the >2 fold upregulated genes in D^high^ (C) and D^low^ (D). E: To validate the wave pattern of gene expression, CXCL10 mRNA was quantified using RT-qPCR in multiple pDC donors (n=3-4) at different time points post TLR7 agonist R837, TLR3 agonist poly(I:C) and SARS-CoV-2 (1 MOI) exposure at indicated time points. Bars represent mean values and equal symbols represent equal donors.

### MyD88 is required for interferon responses to SARS-CoV-2

SARS-CoV-2 - a single stranded RNA virus - may potentially be sensed by the endosomal TLR-MyD88 (Myeloid differentiation primary response 88) pathway (*1, 2, 16*). To evaluate this in detail, we first generated MyD88 knockout (MyD88^KO^) pDCs using CRISPR/Cas9. As a control, we included cells targeted with CRISPR at the inert (safe-harbor) genomic locus AAVS1 (AAVS1^KO^). MyD88 knockout was confirmed by protein expression (Figure 4A), inference of CRISPR edits (ICE) analysis (Figure 4B), as well as lack of type I IFNα and cytokine induction in response to TLR7 agonist stimulation (Appendix Figure S5A-E). Of note, knockout of MyD88 did not affect the pDC phenotype (Appendix Figure S5F). When exposed to SARS-CoV-2, we observed that MyD88^KO^ pDCs were severely impaired in the induction of CXCL10 and type I IFNα, as compared to the AAVS1^KO^ control pDCs (Figure 4C-D).

**Figure 4.**
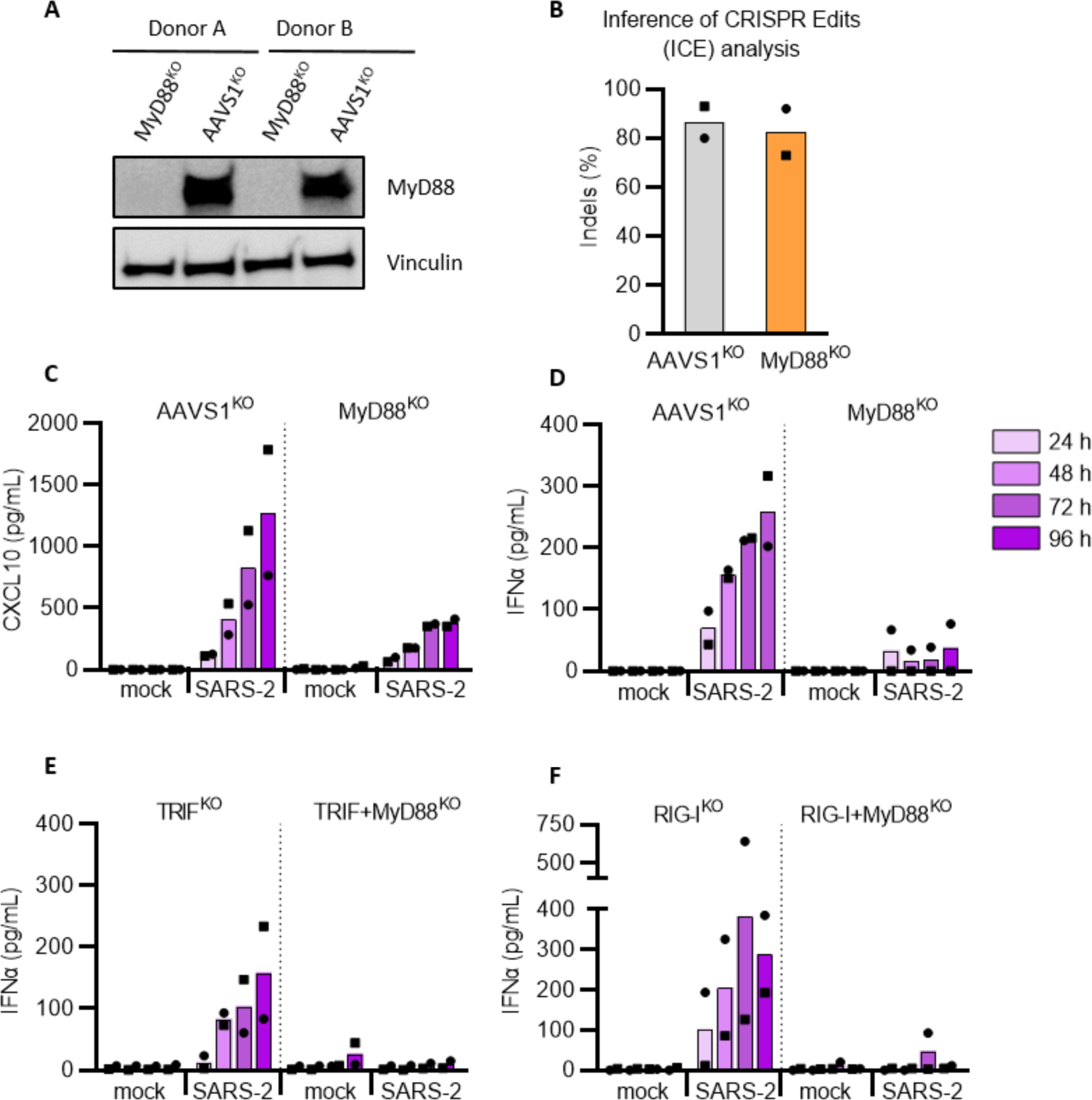
SARS-CoV-2 sensing and inflammatory cytokine induction by pDCs is mediated predominantly via MyD88. Using CRISPR/Cas9, MyD88 knock-out (KO) and AAVS1^KO^ (control) pDCs were generated. MyD88 protein levels in KO and control pDCs were analyzed by western blotting (A) and cellular DNA was sequenced to perform an Inference of CRISPR Edits (ICE) analysis (B). MyD88^KO^ and control pDCs were either mock treated (mock) or exposed to SARS-CoV-2 (SARS-2, 1 MOI), supernatants were collected at indicated time points and analyzed for CXCL10 (C) and type I IFNα (D). Type I IFNα production was then determined in cell culture supernatant from SARS-CoV-2 exposed TRIF^KO^ or TRIG+MyD88^KO^ (E) and RIG-I^KO^ or RIG-I+MyD88^KO^ (F) pDCs. Bars represent mean values and equal symbols represent equal donors (n=2).

Different RNA sensing mechanisms can be active in pDCs and potentially sense SARS-CoV-2; the endosomal TLR3-TRIF (TIR-domain-containing adaptor-inducing IFNβ) cascade and the intracellular RIG-I-MAVS (retinoic acid-inducible gene I/pathway – mitochondrial antiviral signaling protein) (*28-30*). Though TLR3 predominantly binds short double stranded (ds)RNA, it can also bind regions found in secondary RNA structures such as loops and bulges (*31*). As pDCs have been reported to express TLR3, albeit at lower levels than classical myeloid DCs (*29*), we next generated and validated pDCs with a TRIF^KO^ and a double TRIF+MyD88^KO^ (Appendix Figure S6). Disrupting TRIF signaling impaired agonist-induced IFNλ1 production in response to TLR3 agonist (Appendix Figure S6D) but type I IFNα was still produced in response to SARS-CoV-2 exposure (Figure 4E). Next, we tested the RIG-I pathway, which is both expressed and further upregulated in pDCs upon TLR stimulation and type I IFN signaling (*28, 30*). Disrupting RIG-I signaling showed a similar response pattern as observed for the TRIF^KO^ pDCs exposed to SARS-CoV-2, indicating this pathway is not necessary for the sensing of SARS-CoV-2 and subsequent type I IFNα production by pDCs (Figure 4F, Appendix Figure S6). Altogether, our results indicate that pDCs primarily sense SARS-CoV-2 and induce antiviral cytokine production via a MyD88 controlled pathway.

### TLR7 and TLR2 sense SARS-CoV-2 with divergent inflammatory responses

The initial observations of knocking out different signal components in RNA sensing pathways prompted us to narrow down the TLR responsible for sensing SARS-CoV-2 and controlling the induction of cytokines. We first selected to generated pDCs with TLR3^KO^ or TLR7^KO^ pathways. Disrupting these two pattern recognition receptors (Figure 5A-C and Appendix S7) clearly demonstrated that TLR3 was not involved in the production of type I IFNα and CXCL10 post sensing of SARS-CoV-2 (Appendix Figure S7). However, TLR7 knockout completely abolished type I IFNα and showed a trend toward impaired CXCL10 production in response to SARS-CoV-2 exposure, as compared to AAVS1^KO^ control pDCs (Figure 5B-E). Disruption of TLR8, another intracellular viral RNA sensor, with and without TLR7^KO^, confirmed that type I IFNα production in response to SARS-CoV-2 was solely driven by TLR7 (Appendix Figure S8A-D). We also explored the effect of inhibition of the Interleukin 1 Receptor Associated Kinase 4 (IRAK4), as it has previously been shown to be important for SARS-CoV-2-induced cytokine induction in pDCs (*26*). We observed that pDCs treated with IRAK4i prior to viral exposure displayed significantly reduced type I IFNα and CXCL10 protein production, without major effects on cell viability (Figure 5F and S9). Remarkably, we next observed that SARS-CoV-2-induced IL-6 production was completely unaffected by the disruption of the TLR7 and TLR8 sensing pathway (Figure 5G Appendix Figure S8E-F), suggesting a parallel endosomal- and viral RNA-independent sensing mechanism.

**Figure 5.**
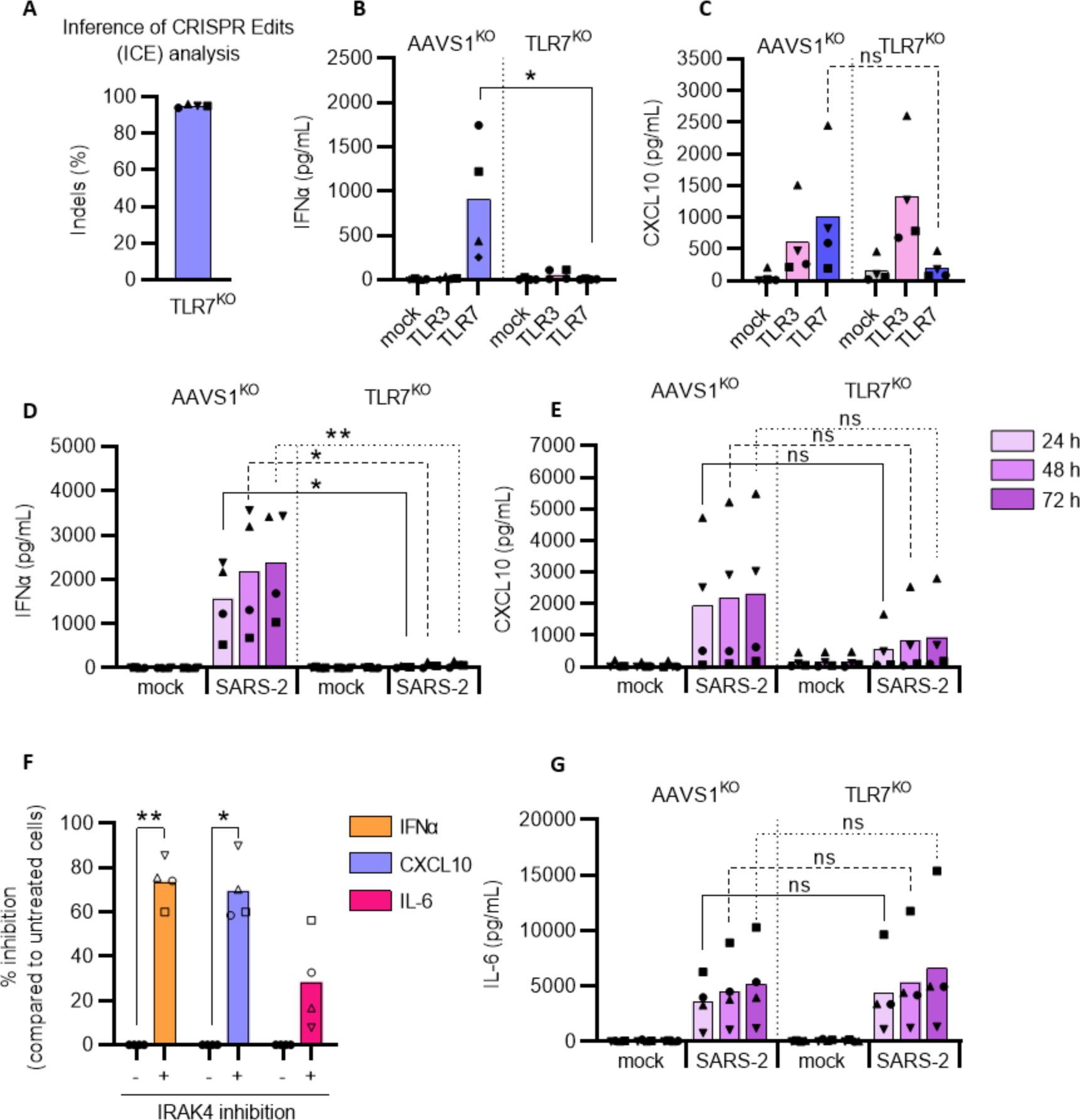
SARS-CoV-2 sensing and subsequent type I IFNα production by pDCs is mediated by TLR7. Using CRISPR/Cas9, TLR7 knock-out (KO) and AAVS1^KO^ (control) pDCs were generated and cellular DNA was sequenced for ICE analysis (A). For functional evaluation each KO pDC donor was stimulated with TLR3 (800 ng/mL poly(I:C, pink) or TLR7 (2.5 μg/mL R837, blue) agonist; supernatant was collected after 24 hrs and analyzed for IFNα (B) and CXCL10 (C) protein expression by ELISA. AAVS1^KO^ and TLR7^KO^ pDCs were either mock treated (mock) or exposed to SARS-CoV-2 (SARS-2, 1 MOI), supernatants were collected at indicated time points and analyzed for type I IFNα (D) and CXCL10 (E) proteins. Wild type pDCs were exposed to SARS-CoV-2 (0.5 MOI) in the absence or presence of an IRAK4 inhibitor (10 μM), 24 hrs after virus exposure the cell culture supernatants were analyzed for production of type I IFNα, CXCL10 and IL-6 proteins (F). IL-6 protein quantification in AAVS1^KO^ and TLR7^KO^ pDCs after SARS-CoV-2 exposure (1 MOI) at indicated time points (G). Bars represent mean values and equal symbols represent the donors used throughout the experiments (n=4). Statistical significance was determined using the ratio paired student T test for agonist or virus treated cells and compared to their respective mock treated conditions, or by unpaired T test when comparing matched conditions between different KOs. *<p0.05, **<p0.01, ns = not significant.

Multiple studies have shown that elevated levels of IL-6 in COVID-19 patients are associated with disease severity (*5, 6, 32, 33*) and thus we next focused on determining what mechanism was responsible for the IL-6 production by pDCs. As murine bone marrow-derived macrophages and human PBMCs can utilize TLR2 to detect SARS-CoV-2 envelope protein (*12*) we hypothesized that this TLR could be engaged by human pDCs to sense SARS-CoV-2 and produce IL-6. First, we generated TLR2^KO^ pDCs (Figure 6A-B and Appendix Figure S10A) and observed that disruption of TLR2 did not affect SARS-CoV-2-mediated type I IFNα production (Figure 6C), but did significantly impair IL-6 production (Figure 6D). Using recombinant glycoproteins of SARS-CoV-2 we next showed that TLR2 sensing and IL6 production was triggered by the envelope protein but not the viral spike protein (Figure 6E). However, neither S or E protein induced the production of type I IFNα (Figure 6F). Importantly, TLR2 is known to form heterodimers with either TLR1 or TLR6 (*34*) suggesting that these receptors could also be involved in SARS-CoV-2 envelope protein sensing. Here, TLR6^KO^ pDCs but not TLR1^KO^ pDCs displayed a disrupted IL-6 production in response to SARS-CoV-2 exposure (Appendix Figure S10), indicating that pDCs produce IL-6 in response to a TLR2 and TLR2/6-mediated sensing of SARS-CoV-2 glycoproteins. These observations were also confirmed in peripheral blood isolated pDCs (Figure 6G-H).

**Figure 6.**
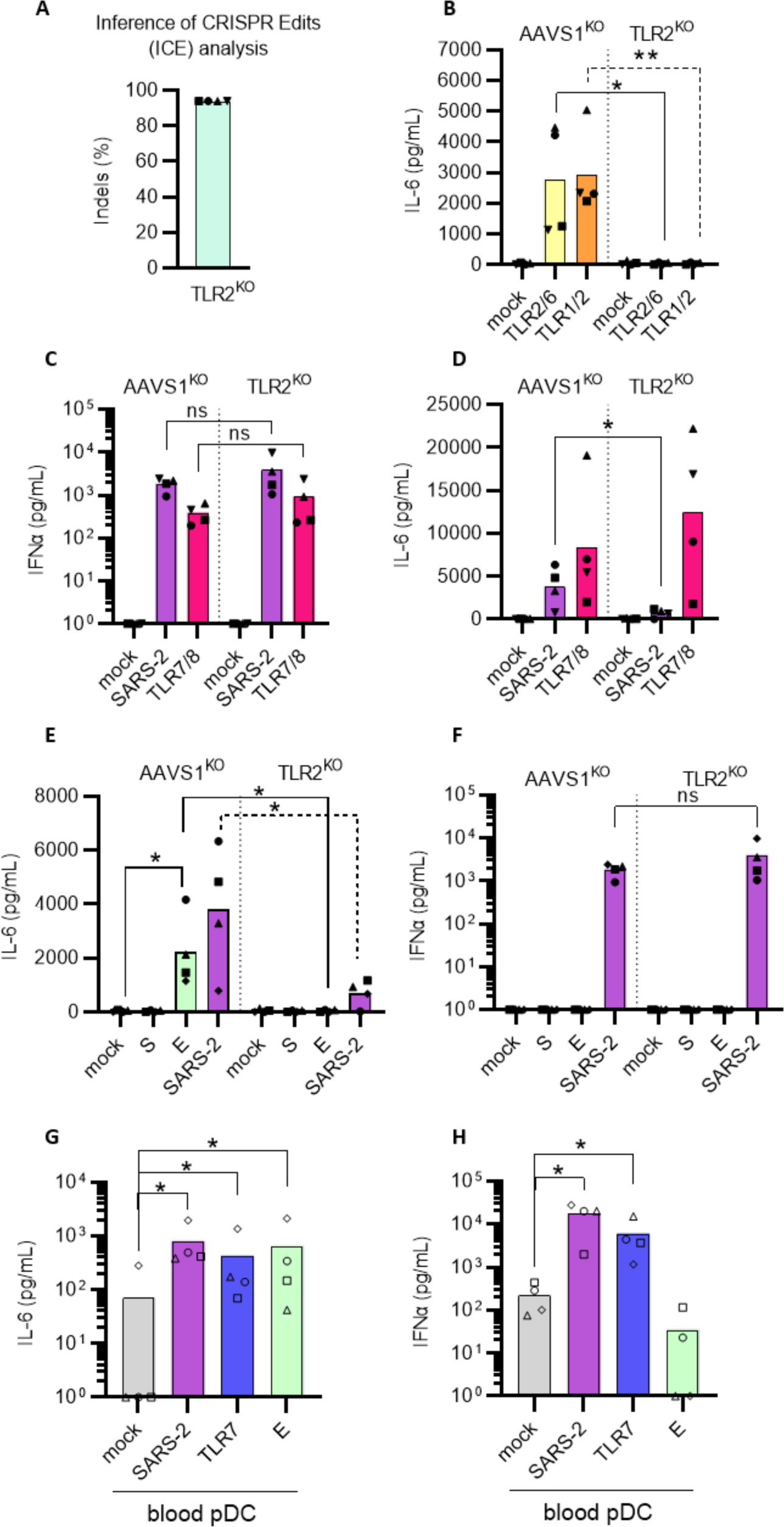
IL-6 production by pDCs is induced by TLR2-mediated sensing of the SARS-CoV-2 envelope protein. Using CRISPR/Cas9, TLR2^KO^ and AAVS1^KO^ (control) pDCs were generated, cellular DNA was sequenced for ICE analysis (A) and cells were evaluated functionally by exposure to two different TLR2 agonists; Pam2CSK4 (5 ng/mL, yellow) and Pam3CSK4 (50 ng/mL, orange) (B). Subsequently, AAVS1^KO^ and TLR2^KO^ pDCs were either mock treated (mock, grey), exposed to SARS-CoV-2 (SARS-2, 0.5 MOI, purple) or TLR7/8 agonist (2.5 μg/mL R848, red) and supernatants were collected after 24 hrs to quantify type I IFNα (C) and IL-6 (D) protein concentrations. To investigate if pDCs sensed the spike or envelope SARS-CoV-2 proteins, AAVS1^KO^ and TLR2^KO^ pDCs were exposed to SARS-CoV-2 (SARS-2, 0.5 MOI, purple) recombinant SARS-CoV-2 spike (S, 1 μg/mL, dark green) or envelope (E, 1 μg/mL, light green) proteins and IL-6 (E) and type I IFNα (F) protein concentrations were quantified. Peripheral blood pDCs were isolated from PBMCs by negative selection and exposed to SARS-CoV-2 (1 MOI, purple), TLR7 agonist (2.5 μg/mL R837, blue) agonist, or E protein (1 μg/mL, light green) for 24 hrs and the concentration of IL-6 (G) and type I IFNα (H) was quantified in cell culture supernatants by ELISA. Bars represent mean values and equal symbols represent equal donors (n=4). Statistical significance was determined using the ratio paired student T test for agonist or virus treated cells and compared to the mock treated condition, or by unpaired T test when comparing matched conditions between different KOs. *<p0.05, **<p0.01, ns = not significant.

### SARS-CoV-2 uses neuropilin-1 to evade the pDCs’ anti-viral response

A few papers have suggested that SARS-CoV-2 can bind to neuropilin-1/CD304/BDCA-4 as alternative to ACE2 for viral entry (*35, 36*). However, ACE2 is not expressed on pDCs (See Appendix Figure S2E-F) (*21, 26*), but interestingly; neuropilin-1 is one of the phenotypic markers for pDCs and often highly expressed on these cells. Neuropilin-1 has been reported to have a functional role in pDCs by reducing type I IFNα production (*37, 38*). Thus, we hypothesize that SARS-CoV-2 engagement with CD304 on the pDCs’ cell surface, may interfere with the immunological responses following viral sensing. To test this, we first evaluated if CD304 receptor engagement, would affect IFNα secretion in our experiments. Exposing pDCs to CD304-specific antibody prior to stimulation with TLR7 agonist, reduced production of IFNα in all donors tested, both stem cell-generated pDCs and blood isolated pDCs (Figure 7A). Next, we generated CD304^KO^ pDCs from multiple donors (Figure 7B). After exposure to SARS-CoV-2 infection, the CD304^KO^ cells showed a strong increase in type I IFNα secretion (up 4.5 fold) at multiple time points (Figure 7C). This clearly indicates that viral engagement with surface neuropilin-1 on the pDC impairs the type I IFNα production by pDCs. Notably, secretion of both pro-inflammatory cytokines CXCL10 and IL-6 production upon SARS-CoV-2 sensing was unaffected in the CD304^KO^ pDCs (Figure 7C-D). This illustrates a novel potential immune evasion strategy of SARS-CoV-2 to reduce the pDCs’ type I antiviral IFNα production without affecting the immunopathological pro-inflammatory responses upon infection.

**Figure 7.**
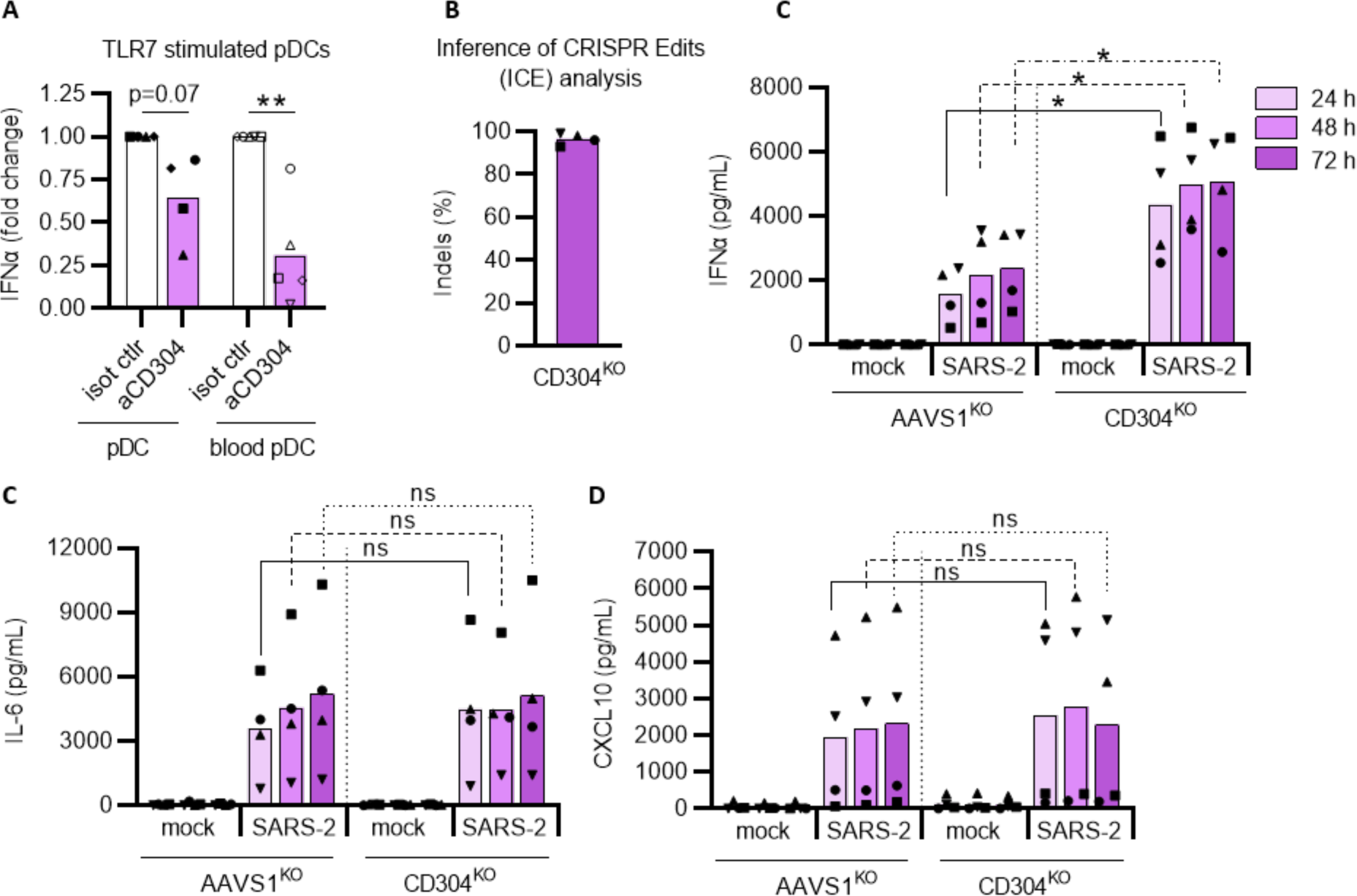
SARS-CoV-2 uses neuropilin-1 as immune evasion strategy by inhibiting type I IFNα production from pDCs. The effect of CD304 signaling on IFNα production was tested by incubating pDCs and peripheral blood isolated pDCs (blood pDC) with anti-CD304 antibody (aCD304) or isotype control antibody (isot ctrl) (10 μg/mL) for 15 minutes prior to stimulation with TLR7 agonist (2.5 μg/mL R837). Cell culture supernatants were harvested after 24 hrs to quantify type I IFNα by ELISA (A). Inference of CRISPR Edits (ICE) analysis of CD304^KO^ pDCs (B). AAVS1^KO^ and CD304^KO^ pDCs were either mock treated (mock) or exposed to SARS-CoV-2 (1 MOI), cell culture supernatants were collected at indicated times and analyzed for type I IFNα (C) IL-6 (D) and CXCL10 (E) protein concentrations. Bars and red lines represent mean values, equal symbols represent equal donors (n=4-5). Statistical significance was determined using the ratio paired student T test for agonist or virus treated cells and compared to the time point-matched mock treated condition, or by unpaired T test when comparing matched conditions between different KOs. *<p0.05, ns = not significant.

## CONCLUSION

Using CRISPR/Cas9-editing of human stem cell-derived pDCs we here demonstrated that pDCs sense SARS-CoV-2 and produce different pro-inflammatory cytokines in response to viral exposure (see graphical abstract). The viral E glycoprotein is recognized by the extracellular

TLR2/6 heterodimer, leading to production of the pro-inflammatory IL-6 cytokine. The intracellular TLR7-MyD88-IRAK4 pathway facilitates the production of CXCL10 and antiviral type I IFNα, of which the latter can protect lung epithelial cells from *de novo* SARS-CoV-2 infection. Removing expression of the NRP1/CD304 receptor from the pDCs’ cell surface, alleviates the SARS-CoV-2-induced inhibition on the antiviral response and enhances type I IFNα production, indicating that SARS-CoV-2 utilizes an intrinsic immune evasion strategy that mitigates antiviral IFN production.

## DISCUSSION

COVID-19 severity is associated with the excessive production of inflammatory cytokines, also described as a ‘cytokine storm’, yet which cells produce these cytokines succeeding SARS-CoV-2 infection is not fully understood. Our findings show that pDCs, an immune cell type important for the host defense against many viruses, efficiently detect SARS-CoV-2 by a multi-faceted sensing mechanism and in response produce inflammatory and antiviral cytokines, including type I IFNα and IL-6.

Since SARS-CoV-2 emerged, multiple studies have suggested that different cell types as well as diverging sensing pathways to be responsible for the control of the viral infection and the increased levels of inflammatory cytokines observed in patients. One of the challenges by exploring the antiviral response of pDCs is the limited number of cells to collect from blood and the notorious difficulties to genetically manipulate these cells. This can partly be overcome by collecting pDCs from patients with genetic disorders (*26*) or by studying mice. However, some TLR pathways have been reported to either being nonfunctional or controversial in mice models (*39, 40*). In the present study, using a stem cell-based human pDC model in combination with CRISPR technology to knockout multiple TLRs and signaling factors, we demonstrated that TLR7 is critical for the inflammatory signal induced by SARS-CoV-2 infection. Unexpectedly, reduction of the inflammatory cytokine IL-6 was solely dependent on the TLR2 pathway whereas TLR7-MyD88 was responsible for the remaining inflammatory cytokines.

Highly pathogenic coronaviruses, similar to other viruses, have multiple strategies to interfere with the host’s immune response and efficient immune evasion is associated with pathogenicity (*23*). Therefore, a detailed understanding of SARS-CoV-2’s immune evasion strategies is critical for the development of antiviral therapeutics. Our data indicate that SARS-CoV-2 utilizes neuropilin-1 not only as alternative receptor to ACE2 for viral entry, but also to mitigate the production of type I IFNα by pDCs, thereby reducing the host’s innate antiviral immune response. The molecular mechanisms leading to the reduce type I IFNα seen when CD304 is present on pDCs will need further investigations.

As pDCs support both the rapid type I IFNα secretion and IL-6 production, this suggests that these cells may have a double-edged function during COVID-19 pathogenesis. Without active pDCs in the lungs, antiviral protection may not be mounted, whereas sustained pDC activation could exacerbate lung inflammation via IL-6 production. Blocking IL-6 responses may not necessarily be successful clinically but therapy with antagonists that specifically impair TLR2, and not TLR7, or therapeutics targeting the viral E glycoprotein could potentially be a scenario to direct immune cells, such as pDCs, to mount a stronger type I and III IFN response that could mitigate disease pathogenesis.

In conclusion, our study provides evidence that circulating pDCs could be a potential therapeutic target to maintain desired antiviral IFN levels allowing for the mitigation of COVID-19 severity.

## METHODS

### HSPC-pDCs

HSPC-pDCs were generated as described previously (*24, 25*). In brief, CD34^+^ HSPCs were purified from human umbilical cord blood (CB) acquired from healthy donors under informed consent from the Department of Gynecology and Obstetrics, Aarhus University Hospital, Aarhus. Mononucleated cells were recovered by standard Ficoll-Hypaque (GE Healthcare) density-gradient centrifugation and CD34^+^ cells were isolated using anti-CD34 immunomagnetic beads (positive selection) following the manufacturer’s instructions (EasySep^TM^ Human cord blood CD34^+^ positive selection kit II, STEMCELL Technologies Cat#17896). CD34^+^ HSPCs were either freshly used or cryo-preserved until future use. For HSPC to pDC differentiation, CD34^+^ HSPCs were cultured using serum free medium SFEM II (STEMCELL Technologies) supplemented with 20 U/mL penicillin and 20 μg/mL streptomycin (Penicillin-Streptomycin ThermoFisher Scientific), 100 ng/mL Flt3-L (Peprotech), 50 ng/mL TPO (Peprotech), 100 ng/mL SCF (Peprotech), 20 ng/mL IL-3 (Peprotech) and 1 µM SR1 (StemCell Technologies). Cells were cultured at 37°C, 95% humidity, and 5% CO2, medium was refreshed every 3-4 days and cells were kept at a density of 0.5-5×10^6^ cells. After a 14-21-day differentiation period, pDCs were enriched using negative magnetic selection, according to the manufacturer’s protocol (EasySep^TM^ Human Plasmacytoid DC Enrichment kit, STEMCELL Technologies Cat#19062). Enriched HSPC-pDCs were then primed for 1-3 days in RF10 (RPMI-1640 medium (Merck) supplemented with 10% (v/v) heat-inactivated fetal calf serum (hiFCS, Sigma-Aldrich), 2 mM L-glutamine (ThermoFisher Scientific), 100 U/mL penicillin, and 100 µg/mL streptomycin) supplemented with 250 U/mL IFNβ (PBL Assay Science), 12.5 ng/mL IFNγ (Peprotech) and 20 ng/mL IL-3. Primed HSPC-pDCs were phenotypically and functionally validated using flow cytometry (Appendix Figure S1B-C) and used for virus inoculation.

#### Cell lines

Calu-3 epithelial lung cancer cells (kindly provided by Laureano de le Vega, Dundee University, Scotland, UK) and human lung adenocarcinoma epithelial A549 cells expressing hACE2 (kindly provided by Brad Rosenberg, Icahn School of Medicine at Mount Sinai, New York, USA) were grown as a monolayer in DMEM10 (Dulbecco’s minimal essential medium, DMEM, Life Technologies, supplemented with 10% (v/v) hiFCS, 2 mM L-glutamine, 100 U/mL penicillin, and 100 µg/mL streptomycin. VeroE6 cells expressing TMPRSS2 (VeroE6-hTMPRSS2, kindly provided by Professor Stefan Pöhlmann, University of Göttingen) (*41*) were grown in DMEM5 (DMEM supplemented with 5% (v/v) hiFCS, 2mM L-glutamine, 100 U/mL penicillin, and 100 µg/mL streptomycin), supplemented with 10 μg/mL blasticidin (Invivogen) to maintain TMPRSS2 expression. All cells were cultured at 37°C and 5% CO2.

#### Air-liquid interface (ALI) epithelium model

ALI cells were generated and cultured as described previously (*10, 42*). In brief, primary nasal cells were isolated using a nasal brush (Dent-O-Care). Cells were cultured as a monolayer in tissue culture flasks coated with 0.1 mg/mL Bovine type I collagen solution (Sigma-Aldrich). At passage two, cells were seeded at 2–3 × 10^~^4 cells on 6.5 mm Transwell membranes (Corning) coated with 30 μg/mL Bovine type I collagen solution and cultured in 2x P/S (200 U/mL Pen/Strep DMEM-low glycose (Sigma-Aldrich) mixed 1:1 (v/v) with 2x Monolayer medium (Airway Epithelium Cell Basal Medium, PromoCell, supplemented with 2 packs of Airway Epithelial Cell Growth

Medium Supplement, PromoCell, without triiodothyronine + 1 mL of 1.5 mg/mL BSA). When cultures reached confluency, Air-liquid interface (ALI) is introduced and medium is changed to ALI medium (Pneumacult ALI medium kit (StemCell) + ALI medium supplement (StemCell) + 100 U/mL Pen/strep) supplemented with 0.48 μg/mL of hydrocortisone (StemCell) and 4 μg/mL heparin (StemCell). Cells were allowed to differentiate for at least 21 days, as verified by extensive cilia beating and mucus covering, prior to experiment initiation.

#### PBMCs and plasma samples from SARS-CoV-2 infected individuals

The study population was derived from a cohort of PCR-confirmed hospitalized COVID-19 patients who were enrolled in a clinical trial (*43*). Individuals for whom there were no cryopreserved peripheral blood mononuclear cells (PBMCs) at baseline, who were pregnant, breastfeeding or had serum total bilirubin x3 above upper limit of normal were excluded from the study. Peripheral blood was collected at time of hospitalization (day 1) and after 5 days. Peripheral blood mononuclear cells (PBMCs) and plasma were separated using ficoll gradient centrifugation, aliquoted, and stored in liquid nitrogen. Self-reported earliest symptom experience was used as determinant for onset of clinical symptoms prior to presentation at the hospital.

#### Isolation of pDCs from peripheral blood for infection studies

PBMCs were isolated by ficoll-Hypaque density centrifugation of standard blood donor buffy coats obtained from Aarhus University Hospital Blood bank. Blood pDCs were enriched using negative magnetic selection, according to the manufacturer’s protocol (EasySep^TM^ Human Plasmacytoid DC Enrichment kit, STEMCELL Technologies Cat#19062). Obtained cells were phenotypically validated by flow cytometry.

### Flow cytometric analysis

Phenotypic validation of HSPC-pDCs and analysis of ACE2 expression was performed using flow cytometry. Briefly, 1-2 × 10^5^ cells were washed with facs wash (FW, PBS supplemented with 1% hiFCS and 0.05 mM EDTA (ThermoFisher Scientific)) and stained in FW with antibodies either 30 min on ice or 15 min at room temperature in the dark. Cells were then washed three times and fixated using 1% formaldehyde (Avantor, VWR, Denmark). Fluorescent intensity was measured with a NovoCyte 3000 Analyzer equipped with three lasers (405, 488, and 640 nm) and 13 PMT detectors (ACEA Biosciences, Inc). Data were analyzed using DeNovoSoftware FCS express flow research edition version 6. OneComp eBeads Compensation Beads (ThermoFisher scientific) were used to compensate for fluorescent spillover and gates were set using fluorescent minus one (FMO) controls in each individual experiment. Cells were gated using the following strategy; total cells (SSC-H/FSC-H); single cells (FSC-A/FSC-H); viable cells (LIVE/DEAD Fixable Near-IR Dead Cell Stain Kit, ThermoFisher Scientific Cat#L10119); negative for lineage markers CD3, CD14, CD16, CD19, CD20, CD56 (anti-human Lineage cocktail 1 – FITC, BD FastImmune Cat#340546) and negative for CD11c (APC mouse anti-human CD11c, clone B-ly6, BD Pharmingen Cat#559877), and subsequently analyzed for the expression of pDC markers CD123 (PE mouse anti-human CD123, clone 6H6, eBioscience Cat#12-1239-42) and CD304 (BV421 anti-human CD304, clone 12C2, BioLegend Cat#354514) (Appendix Figure S1). In some experiments, cells were stained for ACE2 expression (PerCP mouse anti-human ACE2, clone AC384, Novus Biologicals Cat#NBP2-80038PCP).

Quantification of pDCs and mDCs in PBMCs from SARS-CoV-2 infected individuals and healthy controls was performed by flow cytometry. For each time point, 10 million cryopreserved PBMCs were thawed and stained with viability dye for 20 minutes at 4°C, non-specific binding was blocked using Fc Receptor blocking solution (Human TruStain FcX, BioLegend) and cells were stained with antibodies in PBS supplemented with 2% hiFCS for 30 minutes at 4°C. Fluorescent intensity was measured with a BD LSR-Fortessa X-20, using BD FACSDiva Software, gates were set using fluorescent minus one (FMO) controls and data were analyzed with Flow-Jo software. Total dendritic cells (DCs) were gated using the following gating strategy: total single cells (FSC-H/FSC-A followed by SSCA-A/FSC-A); viable cells (Invitrogen LIVE/DEAD Fixable Near-IR Dead Cell Stain Kit); negative for lineage markers CD3, CD14, CD16, CD19, CD20, CD56 (anti-human Lineage cocktail 1 – FITC, BioLegend 348801). From total DCs, myeloid DCs were quantified by gating for CD11+ cells (APC, clone 3,9 TONBO bioscience 20-0116-T100) and pDCs were quantified by gating for CD303+ cells (PE-Cy7, clone 201A 25-9817-42, eBioscience) followed by gating for CD123+ cells (PE, clone 6H6, eBioscience).

### Virus and propagation

The SARS-CoV-2 strain FR2020 was kindly provided by Professor Georg Kochs (University of Freiburg) and Professor Arvind Patel (University of Glasgow, UK) kindly provided the SARS-CoV-2 alpha variant. Virus was propagated using VeroE6 cells expressing human TMPRSS2 (*41*). In brief, 4-6×10^6^ cells were seeded in 5 mL medium in a T75 culture flask and infected at 0.05 multiplicity of infection (MOI). One hour after infection, culture medium was increased up to 10 mL and virus propagation continued up to 72 hrs after infection or if a cytopathic effect (CPE) of approximately 70% was visible. To harvest the virus, cell culture supernatant was removed from the flask, centrifuged at 300g for 5 minutes to remove cell debris, aliquotted and stored at -80°C. The amount of infectious virus in the generated stock was determined using a limiting dilution assay.

### Infection assays

2×10^5^ pDCs were seeded in a 48-well in 100 μL RF10 supplemented with 20 ng/mL IL-3. 100 μL control medium, medium containing SARS-CoV-2 at 1, 0.1, 0.01 MOI, 2.5 μg/mL R837 for TLR7 stimulation (Imiquimod, InvivoGen), 800 ng/mL poly(I:C) for TLR3 stimulation (Poly(I:C) LMW, InvivoGen), 2.5 μg/mL R848 for TLR7 and TLR8 stimulation (Resiquimod, InvivoGen), 2.5 μg/mL CpG-A (ODN2216, Invivogen), 50 ng/mL Pam3CSK4 (Invivogen) for TLR1/2 stimulation or 5 ng/mL Pam2CSK4 (Invivogen) for TLR2/6 stimulation was added for 4 hrs after which the culture was topped up with RF10+IL-3 to a final volume of 1 mL. Cells and supernatants were collected at 4 hrs, 24 hrs, 48 hrs, 72 hrs and 96 hrs post virus inoculation. Supernatants were aliquotted and stored at -80°C until further analysis by ELISA, MSD or limiting dilution assay. Cells were washed with PBS and stored as pellets at -80°C until further analysis by RT-qPCR. In some experiments, the SARS-CoV-2 envelope (E) protein (ABclonal RP01263) or the SARS-CoV-2 spike (S) protein (ABclonal RP01283LQ) was added to pDCs at a final concentration of 1 μg/mL. The IRAK4 inhibitor (Pf06650833, Sigma-Aldrich PZ0327) was used at a final concentration of 10 μM. VeroE6 cells constitutively produce low level IL-6 independently of SARS-CoV-2 propagation. Thus to discriminate between *de novo* IL-6 production by pDCs upon SARS-CoV-2 exposure, mock Vero-virus conditions were run in parallel and the IL-6 signal was subtracted from the actual infection samples, to properly determine IL-6 production by pDCs.

In some experiments, CD304 targeting antibodies were used. Here, pDCs were incubated with anti-CD304 (Purified anti-human CD304 (Neuropilin-1), clone 12C2, BioLegend Cat#354502) or isotype control (Ultra-LEAF Purified mouse IgG2a, clone MOPC-173, BioLegend Cat#400264) antibody for 15 minutes prior to stimulation with TLR7 (2.5 μg/mL R837) agonist for 4 hrs in 200 uL, after which the culture volume was topped up to 1 mL. The final concentration (after topping up the culture volume) of each antibody was 10 μg/mL.

1×10^5^ Calu-3 or A459 hACE2 were seeded in a 48-well in 500 μL DMEM10 and the following day, medium was replaced with 200 μL HSPC-pDC conditioned medium or 200 μL DMEM10. After 18 hrs, cells were inoculated with SARS-CoV-2 at 0.1 MOI, and after 1 hr the cultures were topped up using DMEM10 to a final volume of 1 mL. Supernatants were collected 48 hrs after virus inoculation, aliquoted and stored at -80°C until the viral titers were quantified using a limiting dilution assay. To generate HSPC-pDC conditioned medium, HSPC-pDCs were inoculated at 1 MOI or left unexposed. After 3 days supernatants were stored at -80°C until commencement of the protection experiment. To dilute HSPC-pDC conditioned medium, the medium was diluted 3-fold using DMEM10. To test whether type I IFN contributes to the pDC-mediated inhibition of SARS-CoV-2 inhibition, antibodies blocking the type I IFN receptor (mouse anti-human IFNAR2 antibody, clone MMHAR-2, PBL Assay Science Cat#21385-1) or isotype control (Ultra-LEAF Purified mouse IgG2a, clone MOPC-173, BioLegend Cat#400264) were added to Calu-3 cells in 50 μL PBS and antibodies neutralizing IFNα (mouse anti-human IFN alpha antibody, clone MMHA-2, PBL Assay Science Cat#21100-2) or isotype control (Purified mouse IgG1, clone MOPC-21, BioLegend Cat#400102) were added to 200 μL HSPC-pDC conditioned medium, 10 minutes prior to addition of conditioned medium to the Calu-3 cells. The final concentration (after topping up the culture volume) of each antibody was 10 μg/mL.

### Limiting dilution assay

To determine the amount of infectious virus in cell culture supernatant or generated virus stocks, a limiting dilution assay was performed. 2×10^4^ VeroE6-TMPRRS2 cells were seeded in 50 μL DMEM5 in a 96 well plate. The next day, samples were thawed and 5x diluted, followed by 10-fold serial dilution using DMEM5, and 50 uL of each dilution was added to the cells. Final dilution range covered 10^-1^ – 10^-11^ in quadruplicate for supernatants or octuplicate for virus stocks. Each well was evaluated for cytopathic effect (CPE) by eye using standard microscopy, and the tissue culture infectious dose 50 (TCID50/mL) was calculated using the Reed and Muench method (*44*). To convert to the mean number of plaque forming units (pfu)/mL, the TCID50/mL was multiplied by factor 0.7 (ATCC – Converting TCID[50] to plaque forming units (PFU)). Additionally, cells were fixed by adding 10% Formalin (Sigma-Aldrich) at a 1:1 (v/v) ratio, stained with crystal violet solution (Sigma-Aldrich) and stored at room temperature.

### Reverse transcriptase-quantitative PCR (RT-qPCR)

To determine expression levels of the human *IFNa2a, TNFa, CXCL10, IFNL1, GAPDH, ACE2, TMPRSS2* and SARS-CoV-2 *N1* gene, RNA was purified from cells using the RNeasy mini kit (Qiagen) according to the manufacturer’s instructions with RNA being eluted in 30 μL. Subsequently, 100-200 ng of RNA was used as input for cDNA synthesis using the iScript cDNA synthesis kit (Bio-Rad) on an Arktik thermal cycler (Thermo scientific) with program: 5’25°C; 20’46°C; 1’95°C; 4°C. For commercially available Taqman assays (*IFNa2a* Hs00265051_s1, *CXCL10* Hs00171042_m1, *GAPDH* Hs02758991_g1, *ACE2*: Hs01085333_m1 and *TMPRSS2*: Hs01122322_m1, ThermoFisher), samples were analyzed in a 10 μL (final volume) reaction mix containing; 5 μL Taqman Fast Advanced Master Mix, 0.5 μL Taqman assay, 3.5 μL Nuclease-free water and 1 μL of cDNA. For the SARS-CoV-2 *N1* gene qPCR, primers and probe sequences were provided by the CDC and purchased from Eurofins. Samples were analyzed in a final volume of 10 μL, containing 5 μL Taqman fast Advanced Master Mix, 1 μL fw primer (5 pmol/μL 2019-nCoV-N1 fw primer - GAC CC AAA ATC AGC GAA AT), 1 μL rev primer (5 pmol/μL 2019-nCoV_N1 rev primer – TCT GGT TAC TGC CAG TTG AAT CTG), 1 μL probe (1.25 pmol/μL 2019-nCoV_N1 Probe – FAM-ACC CCG CAT TAC GTT TGG TGG ACC-BHQ1), 2 μL Nuclease-free water and 1 μL of cDNA. Analysis was performed on a Lightcycler 480 platform with program: 2’50°C; 2’95°C; 40x(1”95°C; 20”60°C). Ct values were extracted using the Lightcycler Software.

### Cytokine quantification assays

Supernatants were thawed at room temperature or 4°C and inactivated by adding 1:1 (v/v) 0.4% Triton-X-100. Protein levels were quantified using the Human DuoSet ELISAs for IL-6, IL-8, TNFα, CXCL10 (R&D Systems) or the Human IFN-a pan ELISA kit (Mabtech 3425-1M-6, detecting IFN-a subtypes 1/13, 2, 4, 5, 6, 7, 8, 19, 14, 16 and 17), according to the manufacturer’s instructions on a Synergy SynergyHTX multi-mode platereader (BioTek) using the Gen5 version 3.04 program. Protein levels of IFNα2a, IFNβ, IFNγ and IFNλ1 were quantified using the human U-plex Interferon Combo (Meso Scale Discovery K15094K-2), according to the manufacturer’s instructions on the MESO QuickPlex SQ 120. IL-3, IL-6, IL-8, TNFα and CXCL10 protein levels in plasma from SARS-CoV-2 infected individuals were quantified using the V-PLEX Custom Human Cytokine 54-plex kit (Meso Scale Discovery) according to the manufacturer’s instructions with overnight incubation of the diluted samples and standards at 4°C. For IFNα2a the S-PLEX Human IFNα2a Kit (MSD Cat #K151P3S) was used according to the manufacturer’s instructions. The electrochemiluminescence signal (ECL) was detected by MESO QuickPlex SQ 120 plate reader (MSD) and analyzed with Discovery Workbench Software (v4·0, MSD).

### Generating genetically modified HSPC-pDC

HSPC-pDCs were genetically modified as previously described (*24*). Briefly, sgRNAs directed at *MyD88* (5’-GCTGCTCTCAACATGCGAGTG-3’) (*24*), *TICAM1* (TRIF #1: 5’-AACACATCGCCCTGCGGGTT-3’ and TRIF #2: 5’-CTGGCGACCCCTGTCGCGTG-3’), *DDX58* (RIG-I #1: 5’-GGTGTTGTTTACTAGTGTTG-3’ and RIG-I #2: 5’-GGCATCCCCAACACCAACCG-3’), *TLR1* (TLR1 #1 5’-CAACCAGGAATTGGAATACT-3’ and TLR1 #2 5-CTGATATTCAAATGAGCAAT-3’), *TLR2* (TLR2 #1 5’- CTAAATGTTCAAGACTGCCC-3’ and TLR2 #2 5’- AATCCTGAGAGTGGGAAATA-3’), *TLR3* (TLR3 #1: 5’-GTACCTGAGTCAACTTCAGG-3’ and TLR3 #2: 5’- CTGGCTATACCTTGTGAAGT-3’), *TLR6* (TLR6 #1 5’-TTCCAACTATTATGATCATA-3’ and TLR6 #2 5’- CAAGTAGCTGGATTCTGTTA-3’), *TLR7* (TLR7 #1: 5’- CTGTGCAGTCCACGATCACA-3’ and TLR7 #2: 5’- TCCAGTCTGTGAAAGGACGC-3’), *TLR8* (TLR8 #1 5’-GTGCAGCAATCGTCGACTAC-3’ and TLR8 #2 5’- TCCGTTCTGGTGCTGTACAT-3’), CD304/NRP-1 (CD304 #1 5’- CCCGGGTACCTTACATCTCC-3’ and CD304 #2 5’-CTGTCCTCCAAATCGAAGTG-3’) and *AAVS1* (control sgRNA, 5’-GGGGCCACTAGGGACAGGAT-3’) (*24*) were synthesized by Synthego with the three terminal nucleotides in both ends chemically modified with 2′-*O*-methyl- 3′-phosphorothioate. Thawed CD34^+^ HSPCs were cultured at low density (10^5^ cells/mL) for 3-4 days in SFEM II medium supplemented with 20 U/mL penicillin and 20 µg/mL streptomycin, 100 ng/mL Flt3-L, 50 ng/mL TPO, 100 ng/mL SCF, and 35 nM UM171 (STEMCELL technologies). Ribonucleoprotein (RNP) complexes were made by incubating 6 μg Cas9 protein (Alt-R S.p.Cas9 Nuclease V3, Integrated DNA Technologies) with 3.2 μg sgRNA in a final volume of 2 μL at room temperature for 15-20 minutes. 200.000 – 800.000 HSPCs were washed with PBS, resuspended in 20 μL 50 mM Mannitol buffer (made in house; 5 mM KCl, 120 mM Na2HPO4/NaH2PO4, pH 7.2, 15 mM MgCl2), added to the RNP complexes and transferred to a Nucleocuvette strip chamber (Lonza). In case of multiple sgRNAs were used for nucleoporation, individual sgRNAs were incubated with Cas9 protein, after which they were pooled and added to the cells. Nucleoporation was performed using the Lonza 4D-Nucleofector^TM^ System (program DZ100) and HSPC were subsequently cultured for 21 days in HSPC-pDC differentiation medium as described above. CRISPR-Cas9 induced genetic modification were validated at the genomic and protein level, as described below.

### DNA extraction, PCR and sequencing

To validate the CRISPR-Cas9 induced genetic modification at the genetic level, cells were harvested and DNA was extracted using the DNeasy Blood & Tissue Kit (Qiagen Cat#69504). Amplicons were generated using 100 ng DNA as input in a final volume of 40 μL (consisting of 8 μl 5x Phusion GC buffer (Phusion High-Fidelity DNA Polymerase set (ThermoFisher Scientific, Cat#F534), 0.8 μL dNTPs (dNTP Set, 100 mM, InvivoGen Cat#10297117), 0.4 μL Phusion Green High-Fidelity DNA polymerase (ThermoFisher Scientific), 2 μL (10 μM) fw primer, 2 μL (10 μM) rev primer and nuclease free water) on an Arktik Thermal Cycler (ThermoFisher Scientific) with program: 1’98°C; 35x(10”98°C; 30”68°C;1’72 °C; 10’72 °C; 4 °C). The following primers were used: MyD88 fw: 5’-CTC CGT GGA AGA ACT GTG GC-3’; MyD88 rev: 5’-GGC GGC TGT ATC CAA CGC-3’; AAVS1 fw: 5’-TCA GTG AAA CGC ACC AGA CA–3’; AAVS1 rev 5’- CCA CTA CTA CGC CTG GAT GT-3’; TRIF fw: 5’-AAA CCA GCA CCA ACT ACC CA-3’; TRIF rev #1: 5’-TAG GCT GAG TAG GCT GCG TT-3’; TRIF rev #2: 5’-CCC CCA AAG GGC ATT CGA G-3’; RIG-I fw #1: 5’-CTA AGG ACT TGC CTA CAG CT-3’; RIG-I fw #2 5’-GGC TCT GTG CTA AGG ACT TG-3’; RIG-I rev #1: 5-TGC TTG GGA TGA GAG CTC AG-3’; RIG-I rev #2: 5’-CAG ATA GCC AAG AGC TGG GC-3’; TLR1 fw: 5’-TGG TGA GCC ACC ATT CAA CC-3’; TLR1 rev: 5’-TGC GTG TAC CAG ACA CTG TG-3’; TLR2 fw: 5’-CTT GCT CTG TAA TTCC GGA TGG-3’; TLR2 rev: 5’-TGC AGC CTC CGG ATT GTT AAC-3’; TLR3 fw: 5’-AGC TGC AAC TGG CAT TAG GGT G-3’; TLR3 rev: 5’-GGG AGA AAG CGA GAG AGG CA-3’; TLR6 fw: 5’-GCC TAT ATT GCC CCT TCT GGC-3’; TLR6 rev: 5’-CCA CAG GTT TGG GCC AAA GA-3’; TLR7 fw: 5’-ATG CTG CTT CTA CCC TCT CGA-3’; TLR7 rev: 5’- AGT AGG GAC GGC TGT GAC AT-3’; TLR8 fw: 5’-TTG GGA TTA CAG GTG TGA GCC-3’; TLR8 rev: 5’-TTG GGA TTA CAG GTG AGC C-3’; CD304 fw 5’- GAA GCT CCC AGG GGA CCA T-3’; CD304 rev 5’- ACA ACA CAA GGG GTC GAA CAG-3’. Amplicons were separated on a 1% agarose gel using FastDigest Green Buffer (10x, ThermoFisher Scientific, Cat#B72), appropriate bands were excised and purified using the E.Z.N.A Extraction Kit (Omega Bio-Tek, Cat#D2500-01), according to manufactures’ instructions. Isolated amplicons (60 ng) were sent for sequencing with 2.5 μM of a single primer in a total volume of 10 μL to Eurofins Genomics. Sequences were subsequently analyzed using the Interference of CRISPR Edits online tool (ICE, Synthego). In addition to validating the mutations, all control samples were validated to have an intact sequence spanning 300 bp up and downstream the targeted region.

### Western Blot analysis

Cells were washed with ice cold PBS and lysed in (200.000 cells/ 100 μL) Ripa buffer (Thermofisher Scientific) supplemented with Pierce protease and phosphatase inhibitors (Thermofisher Scientific, A32961), Complete Ultra protease inhibitor (Roche, 05892791001), Sodium fluoride (Avantar) and Benzonase Nuclease (Sigma-Aldrich), for 15 minutes on ice and stored at -20°C. Samples were thawed on ice, diluted 1:1 (v/v) with Laemmli sample buffer (Sigma-Aldrich, S3401), incubated at 95°C for 4 min, cooled on ice for 5 min, 30 μL was loaded for MyD88 and RIG-I analysis and 40 μL for TRIF analysis, together with Precision Plus Protein Kaleidoscope protein marker (Bio-Rad 1610395) onto a 10% Criterion TGX Precast Midi Protein Gel (18 well Bio-Rad, 5671034) in Nu PAGE MOPS SDS running buffer (Thermo Scientific NP0001). Proteins were transferred onto a Trans-Blot Turbo Midi PVDF Transfer membrane (Bio-Rad, 170-4157) using the Trans-Blot Turbo Transfer System (Bio-Rad). Membranes were washed using Tris-buffered saline (Fisher Scientific) supplemented with 0,05% (v/v) Tween 20 (Sigma-Aldrich) (TBS-T), blocked for 1 hr at room temperature (RT) in 5% Skim Milk Powder (Sigma-Aldrich) in TBS-T, washed with TBS-T, incubated over-night (o/n) at 4°C with primary antibody diluted 1:500 in 5% Bovine Serum Albumin Fraction V (Roche 10735086001) in TBS-T. The following morning, membranes were washed with TBS-T, incubated for 1 hr with secondary antibody diluted 1:7500 in 5% Skim Milk, washed, and proteins were visualized using Clarity Western ECL Substrate (Bio-Rad, 170560) for MyD88 analysis and SuperSignal West Femto Maximum Sensitivity Substrate (ThermoFisher Scientific, 34095) for RIG-I and TRIF analysis on an ImageQuant LAS 4000 mini biomolecular imager (GE Healthcare). Membranes were washed with TBS-T, and incubated o/n at 4°C with the primary antibody diluted 1:10.000 for the loading control. To detect MyD88, rabbit-anti-human (r-a-h)MyD88 (clone D80F5, 33 kDa, Cell Signaling Technology, cat#4283) was used. For TRIF, r-a-hTRIF (98 kDa, Cell Signaling Technology, cat#4596) was used, and for RIG-I, r-a-hRIG-I (clone D14G6, 102 kDa, Cell Signaling Technology, cat#3743) was used. Each membrane was re-used to for the loading control vinculin (mouse-a-hVCL, clone hVIN-1, 116 kDa, Sigma-Aldrich, cat#V9131). As secondary antibodies, peroxidase-conjugated donkey-anti-rabbit and donkey-anti-mouse was used (Jackson Immuno Research 711-036-152 and 715-036-150).

### NanoString nCounter analyses

To perform broad transcriptomic profiling on SARS-CoV-2 exposed HSPC-pDCs, an nCounter NanoString analysis was performed (NanoString Technologies, Seattle, WA, USA). HSPC-pDCs from two donors were exposed to SARS-CoV-2 at 1 MOI (for 4, 24 or 48 hrs and mock treated samples at 4 and 48 hrs), after which cell pellets were collected and RNA extracted with the RNeasy mini kit (Qiagen). 30 ng of RNA was used as input for the analysis using the nCounter SPRINT Profiler (NanoString Technologies) and the nCounter PanCancer Immune Profiling Panel (cat# XT-CSO-HIP1-12) plus a custom made PanelPlus of the following genes: *NFE2L2, TMEM173, MB21D1, IFNLR1, IRF9, IFNL3, IFNL4, AIM2, TREX1, ENPP1, PCBP1, PQBP1, G3BP1, STIM1, LRRC8A, SLC19A1, NLRC3, NLRX1, ZDHHC1, TRIM56, TRIM32, RNF5, ULK1, TTLL4, TTLL6, AGBL5, AGBL4, PRKDC, DDX41.* Analysis was performed according to the manufacturer’s protocol using a 20 hours hybridization time.

The raw data were processed using the nSOLVER 4.0 software (NanoString Technologies) for D^high^ and D^low^ separately to ensure proper normalization of each dataset. Firstly, a positive control normalization was performed using the geometric mean of all positive controls except for the control named F, as recommended by the manufacturer. Finally, a second normalization was performed using the geometric mean of housekeeping genes with reasonable expression levels and low coefficient of variance percentage (*ABCF1*, *AMMECR1L*, *CNOT10*, *CNOT4*, *DDX50*, *EDC3*, *POLR2A*, *TBP*, *TLK2* and *ZNF143* for D^high^ and *G6PD*, *GPATCH3*, *MRPS5*, *MTMR14*, *POLR2A* and *SDHA* for D^low^), before exporting the data to Excel (Microsoft Corporation, Redmond, WA, USA). Background threshold levels were calculated based on the mean plus two standard deviations of the eight negative controls. Genes with an average expression below the threshold were excluded from further analyses. Data were plotted using Prism 8.2.0 (GraphPad, La Jolla, CA, USA) and R software version 3.5.1 with the following packages installed: ggplot2, circlize, dendextend, ComplexHeatmap and RColorBrewer.

### Reactome pathway overrepresentation analysis

To assign pathways to the gene clusters identified in pDCs from D^high^ and D^low^ 48 hrs after SARS-CoV-2 exposure using unsupervised hierarchical cluster analysis on the NanoString nCounter data, we utilized the Reactome Pathway Browser version 3.7, database release 75 (https://reactome.org/PathwayBrowser); a comprehensive web-based resource for curated human pathways. Disease pathways were excluded from the analyses and we used UniProt as the source of entities (maximum pathway size was 400). Only six genes were not assigned to any pathways in Reactome. Reactome defines statistically significantly enriched pathways using a Binomial Test, followed by correction for multiple comparisons by the Benjamini–Hochberg approach (*45*).

### Statistical analysis

Differences between experimental conditions were analyzed using the ratio paired student T test with GraphPad Prism (Version 6). P-values ≤0.05 were considered significant: *p<0.05, **p<0.01, ***p<0.001. To determine correlation between IFNα production by pDCs and time of exposure to SARS-CoV-2, as well as to compare gene expression changes in D^high^ and D^low^ after 4 and 48 hrs after exposure to SARS-CoV-2, and determine correlation between pDC frequency and disease severity, simple linear regression analysis were performed using GraphPad Prism. The R squared and p-value are indicated in the figures.

## Acknowledgments

We thank Laureano de le Vega (Dundee University, Scotland) for providing the Calu-3 cells, Brad Rosenberg (Icahn School of Medicine at Mount Sinai, New York, USA) for providing the A549 hACE2 cells, Professor Stefan Pöhlmann (University of Göttingen, Germany) for providing the VeroE6-hTMPRSS2 cells, Professor Georg Kochs (University of Freiburg, Germany) for providing the SARS-CoV-2 FR2020 strain and Professor Arvind Patel (University of Glasgow, UK) for providing the SARS-CoV-2 B.1.1.7 strain. We thank Hai Qing Tang, Lars Henning Pedersen and Niels Uldbjerg (Department of Obstetrics and Gynaecology, Aarhus University Hospital Skejby, Denmark) for providing cord blood donors. Flow Cytometry was performed at the FACS Core Facility, Aarhus University, Denmark.

## Funding

This project has received funding from the European Union’s Horizon 2020 research and innovation program under the Marie Skłodowska-Curie grant agreement No 754513 and The Aarhus University Research Foundation (RMS); Independent Research Fund Denmark 0214-00001B (SRP); European Research Council (ERC-AdG ENVISION; 786602); Lundbeck Foundation grant R349-2020-419 (OSS), The Central Denmark Region grant A3233(OSS), the Lundbeck Foundation R238-2016-2708 (MRJ) and Independent Research Fund Denmark 8020-00210B (MRJ).

## Author contributions

RMS and MRJ conceptualized the project, and designed the experimental setups. RMS performed most of the experiments with support from AGO, KRG, MI, JJJ, EC, SHG, SDN, JT, TWB, SSH and AL. KRG designed primers for the RT-qPCR, JJJ performed the qPCR, SHG validated the CRISPR/Cas9 edited cells and JGP conducted the NanoString experiment. UA and LSK performed formal analysis on the NanoString data. LBC performed flow cytometry on the patient samples. LR, MI, DO, ROB, JDG, CKH, SRP, MT and OSS provided invaluable resources and support of SARS-CoV-2 infection studies. RMS drafted the manuscript and MRJ reviewed and edited the manuscript. All authors reviewed, edited and approved the final manuscript

## Competing interests

MRJ, ROB and AL are founders and shareholders of UNIKUM Therapeutics that holds license to the commercialization of the pDC technology for cell therapy. The remaining authors declare no competing interests.

## Data and materials availability

All data are available in the main text or the supplementary materials.

## Expanded View figures

**Figure EV1.**
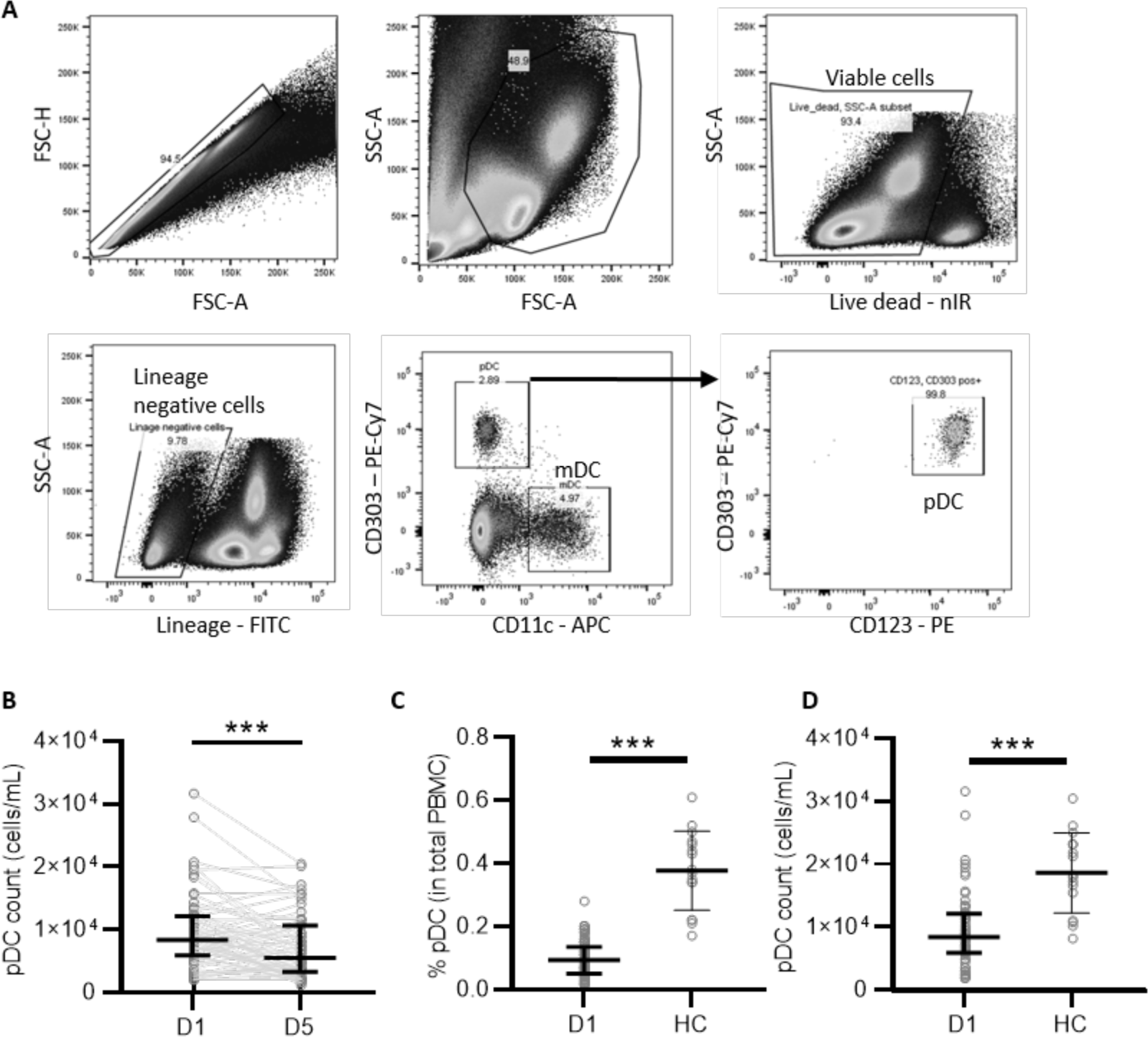
SARS-CoV-2 infection reduces pDC frequency in peripheral blood. A: Overview of the flow cytometric gating strategy to determine the percentage of mDC and pDC from total PBMCs obtained from COVID-19 patients. B: PDC counts / mL of blood collected at time of hospitalization (D1) and 5 days thereafter (D5) (n=62). Percentage of pDC from total PBMC (C) and pDC counts / mL blood (D) from COVID-19 patients at D1 of hospitalization (n=93 and 62, respectively) compared to healthy controls (HC, n=16). Each dot represents a patient, line with error bar shows median value with interquartile range. Statistical significance was determined with the Wilcoxon matched pairs signed rank test. ***<p0.001.

**Figure EV2.**
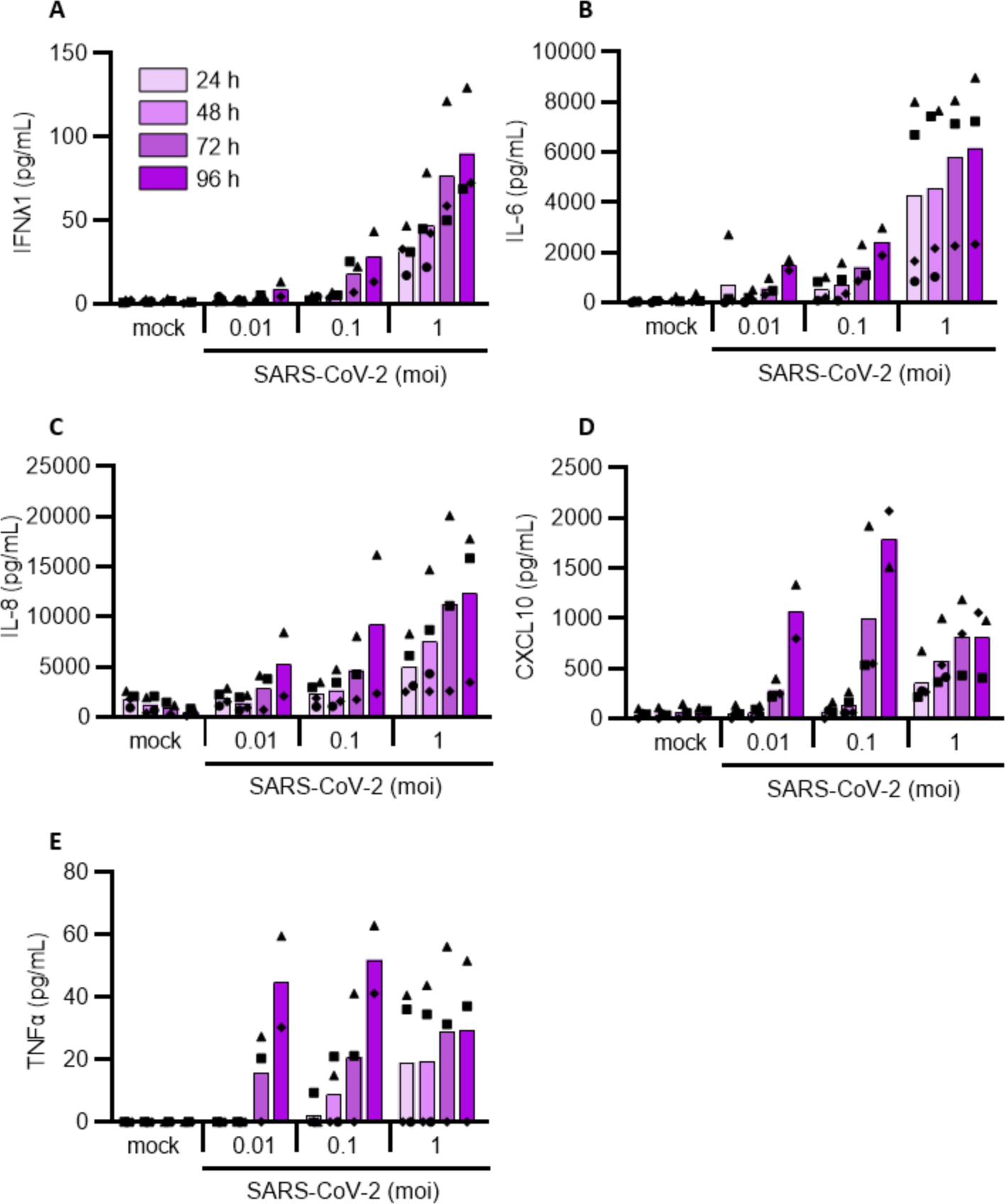
Induction of inflammatory cytokines enhances with increasing viral titers and duration of exposure. HSPC-pDCs were either mock treated (mock) or inoculated with increasing SARS-CoV-2 titers (MOI of 0.01, 0.1 or 1). Supernatant was collected after 24h, 48h, 72h and 96h for the quantification of type III IFNλ1 (A), IL-6 (B), IL-8 (C), CXCL10 (D) and TNFα (E) proteins. Bars represent mean values, symbols represent individual HSPC-pDC donors (n=2-4). Equal symbols represent equal donors.

**Figure EV3.**
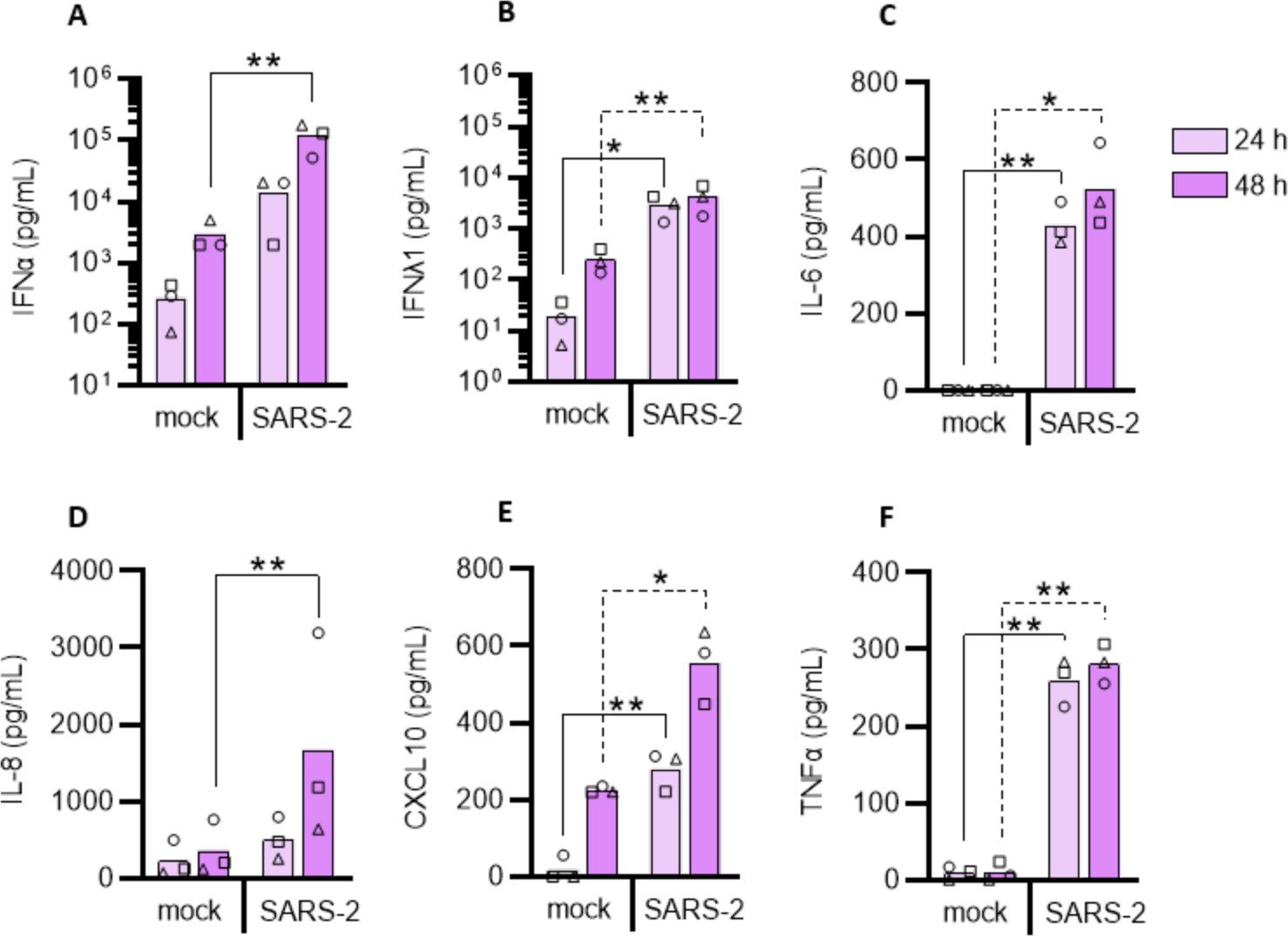
PDCs isolated from peripheral blood sense SARS-CoV-2 and produce inflammatory cytokines similar to stem cell generated pDCs. pDCs were isolated from PBMCs using negative selection and either mock treated (mock) or exposed to SARS-CoV-2 (1 MOI, SARS-2). Supernatants were collected at indicated time points and the production of type I IFNα (A), type III IFNλ1 (B), IL-6 (C), IL-8 (D), CXCL10 (E) and TNFα (F) was quantified by ELISA. Bars and lines represent mean values and symbols represent individual donors (n=3). Statistical significance was determined using the ratio paired student T test by comparing virus exposed cell culture supernatant to the mock control of the matched time point. *<p0.05, **<p0.01.

**Figure EV4.**
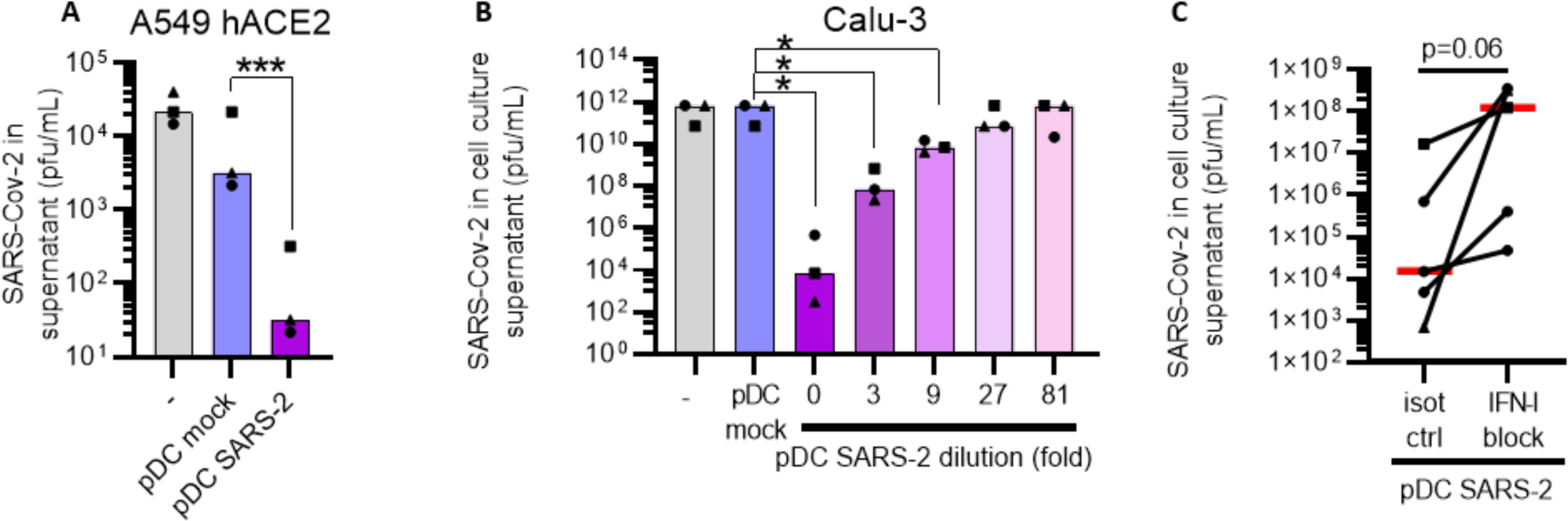
Plasmacytoid DC-secreted cytokines produced post SARS-CoV-2 sensing, protect epithelial cells from infection. To assess whether pDCs could mount protection against SARS-CoV-2, conditioned medium from SARS-CoV-2-exposed pDC cultures (d3 post inoculation with 1 MOI) was added to A549 hACE2 lung epithelial cells (A) or Calu-3 (B) cultures followed by SARS-CoV-2 inoculation. The cell cultures were conditioned with normal medium ( -, grey), pDC supernatant (pDC mock, blue) or SARS-CoV-2-inoculated pDC supernatant (pDC SARS-2, purple), prior to infection with SARS-CoV-2 (0.1 MOI). Supernatants were collected and viral outgrowth was determined 48h post infection. To investigate a potential dose-response, SARS-CoV-2-inoculated pDC supernatant was 3-fold serially diluted prior to addition to Calu-3 cells (purple-pink gradient, B). To determine the involvement of type I IFNs, Calu-3 cells and SARS-CoV-2-inoculated pDC supernatants were pre-treated with antibodies blocking the type I IFN receptor and antibodies neutralizing type I IFNα (IFN-I block) or isotype control antibodies (isot ctrl), prior to the addition of conditioned medium to the cells and infection (C). All antibodies were used at a final concentration of 10 μg/mL. Bars and lines represent median values and symbols represent individual HSPC-pDC donors (n=3-5) Equal symbols represent equal donors. Statistical significance was determined using the ratio paired student T test. *<p0.05, ***<p0.001.

**Figure EV5.**
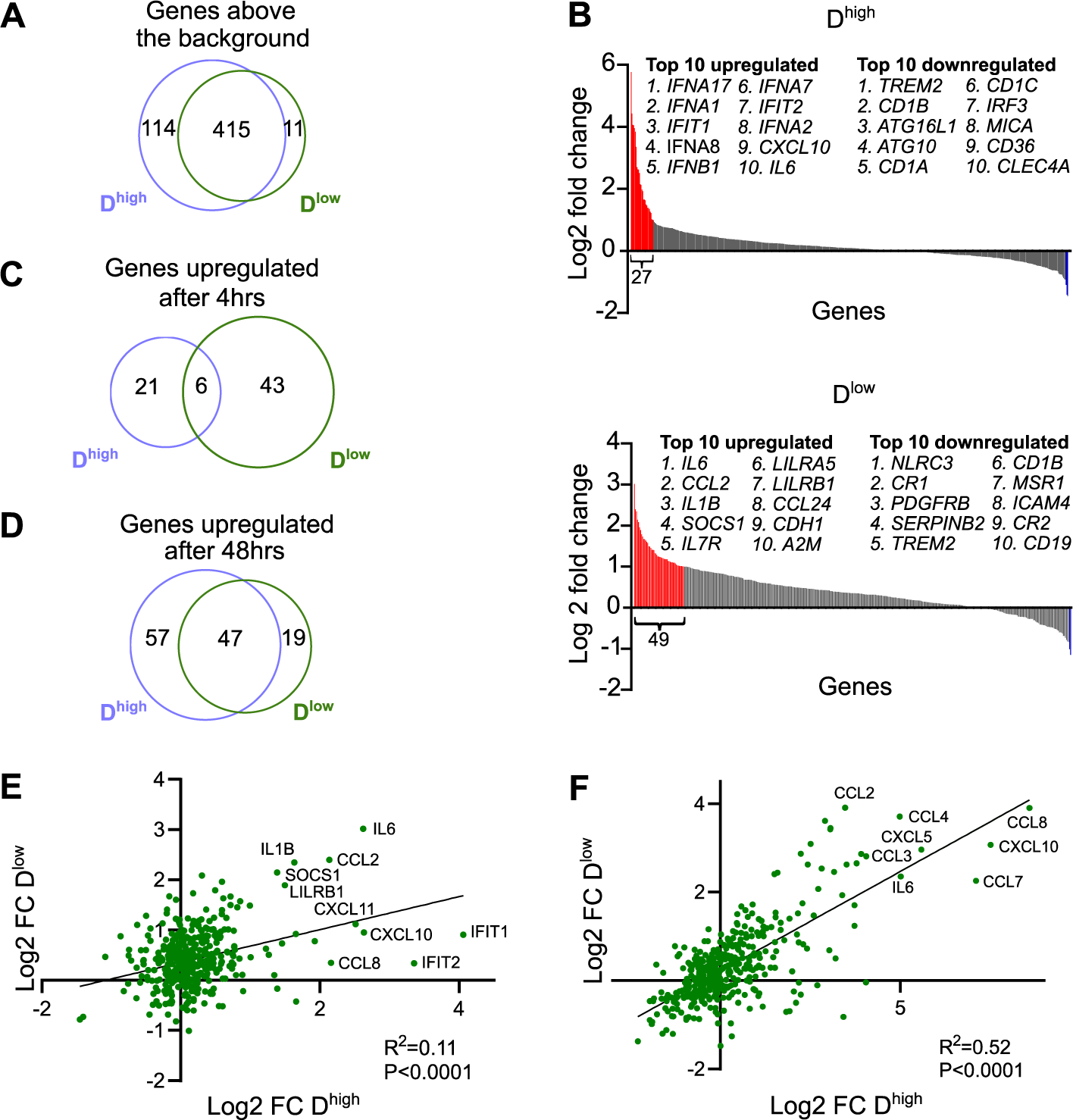
SARS-CoV-2-induced gene expression changes in pDCs. Venn diagram illustrating the large overlap in gene expression detected above background levels in D^high^ and D^low^, measured using a NanoString nCounter (A). Waterfall plots illustrating gene expression changes in pDCs 4h after SARS-CoV-2 exposure relative to mock treated cells from two donors; D^high^ (B, top) and D^low^ (B, bottom). Venn diagrams illustrating the number of genes that were >2 fold upregulated after 4h (C) and 48h (D) in D^high^ and D^low^ post SARS-CoV-2 exposure. Scatter plots of log2 fold changes (FC) in gene expression in D^high^ and D^low^ after 4h (E) and 48h (F) post SARS-CoV-2 inoculation with corresponding linear regression statistics and R-squared values. Note that the genes, which were not detected above background levels in any of the donors, could not be displayed due to division by zero.

## Appendix Supplementary Materials

Appendix Figures S1-S10 Tables S1-S3

