## Supplementary figures for "Distinct SARS-CoV-2 sensing pathways in pDCs driving TLR7-antiviral vs. TLR2-immunopathological responses in COVID-19"

**This PDF file includes:**

Figures S1 to S10
Captions for Tables S1 to S3

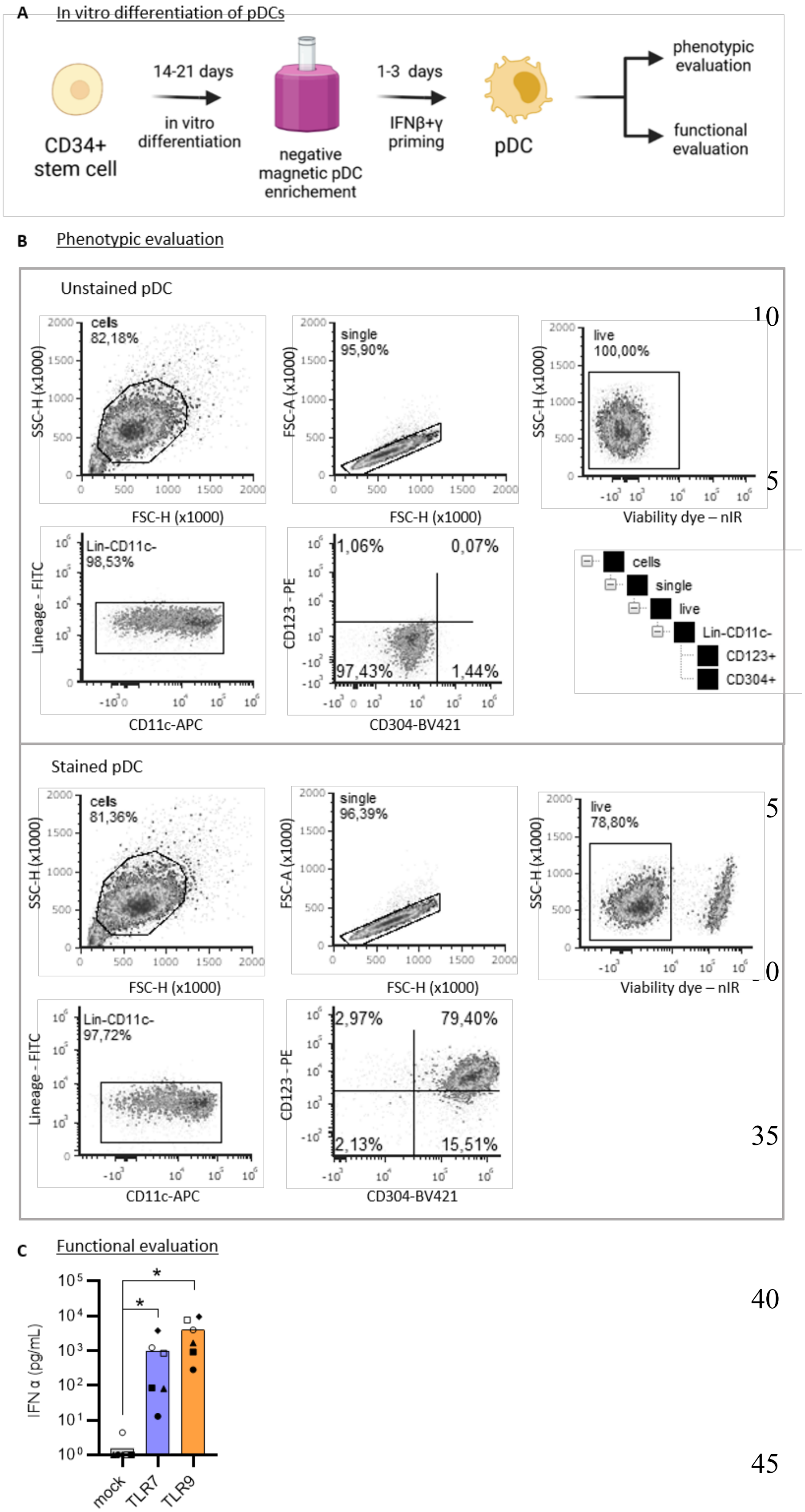


**Figure S1.**

**Generating pDCs from stem cells.** A: CD34+ hematopoietic stem and progenitor cells (CD34+ stem cell) are isolated from human umbilical cord blood and undergo a 14-21 day differentiation period. PDCs are enriched using negative magnetic selection and primed for maturation with IFNbeta (IFNβ) and IFNgamma (IFNγ) for 1-3 days. Each donor is phenotypically (B) and functionally (C) evaluated. B: Overview of the flow cytometric gating strategy to determine the pDCs' phenotype characterized by being Lineage and CD11c negative and positive for CD123 and CD304. Shown are unstained (top) and stained (bottom) pDCs, while gates were set using fluorescent minus one controls. C: Functional evaluation of pDCs, characterized by producing type I IFNα in response to 24 hrs of stimulation with TLR7 (2.5 µg/mL R837) or TLR9 (2.5 µg/mL CpG-A) agonist, as measured by ELISA (n=6). Illustration was created with BioRender.com.

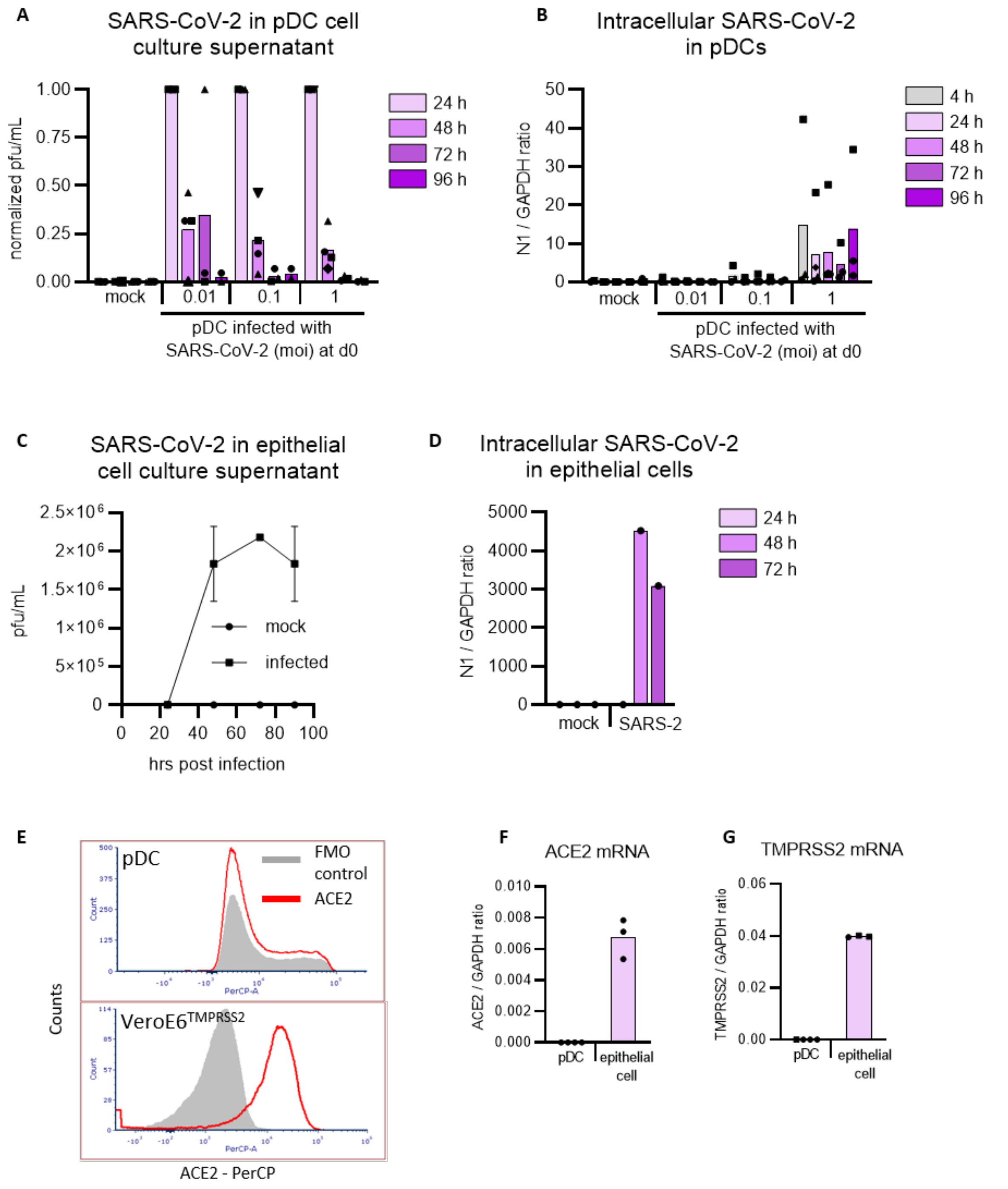


**Figure S2.**

**Plasmacytoid DCs are not a reservoir for SARS-CoV-2 replication.** To assess whether SARS-CoV-2 replicates in pDCs, HSPC-pDCs were mock treated (mock) or exposed to increasing SARS-CoV-2 titers (MOI of 0.01, 0.1 or 1). Supernatants (A) and cells (B) were collected at different time intervals, as indicated, and viral outgrowth in supernatant (A) and intracellular viral amplification (B) was analyzed using the limiting dilution assay and qPCR, respectively. Virus in cell culture supernatant was normalized to viral titers at the 24 hr time point. To confirm that the utilized assays detected SARS-CoV-2 replication, air liquid interface (ALI) epithelial cell cultures were used as a control for productive infection of SARS-CoV-2 (0.5 MOI) and viral outgrowth (C) as well as intracellular SARS-CoV-2 amplification (D). ACE2 expression was measured on pDC and VeroE6 TMPRSS2 by flow cytometry (E). Quantification of *ACE2* (F) and *TMPRSS2* (G) mRNA in pDC and epithelial cells (not detectable was set to 0). Bars represent mean values, symbols represent individual pDC donors (n=3-4).

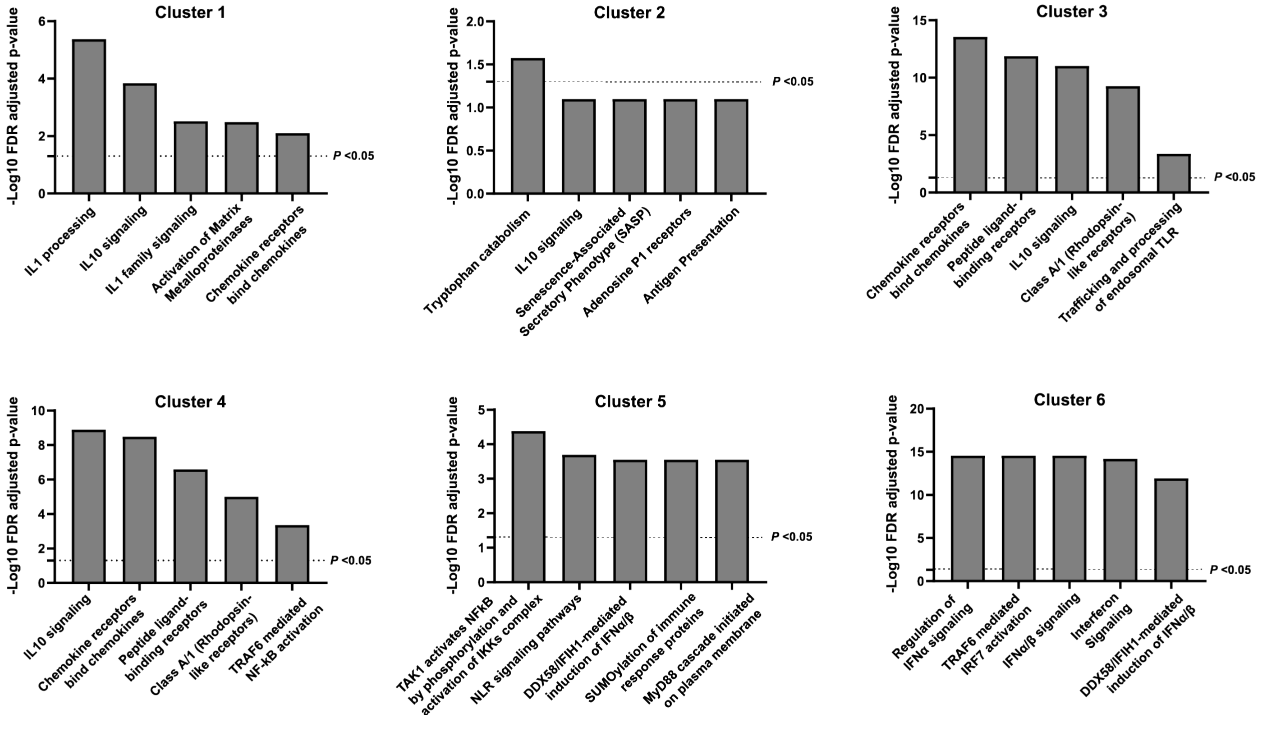


**Figure S3.**

**Reactome Pathway Analysis for donor D$^{high}$.** Histograms illustrating the five most significant pathways and the associated False Discovery Rate (FDR) corrected P-values for the gene clusters derived from the unsupervised hierarchical clustering analysis displayed in Fig 4C for D$^{high}$. Reactome uses the Binomial Test to assess the probability that the overlap between the query and the pathways has occurred by chance and the FDR is calculated using the Benjamini-Hochberg approach. Dotted lines indicate statistical significance. Antigen representation: Folding, assembly and peptide loading of class I MHC.

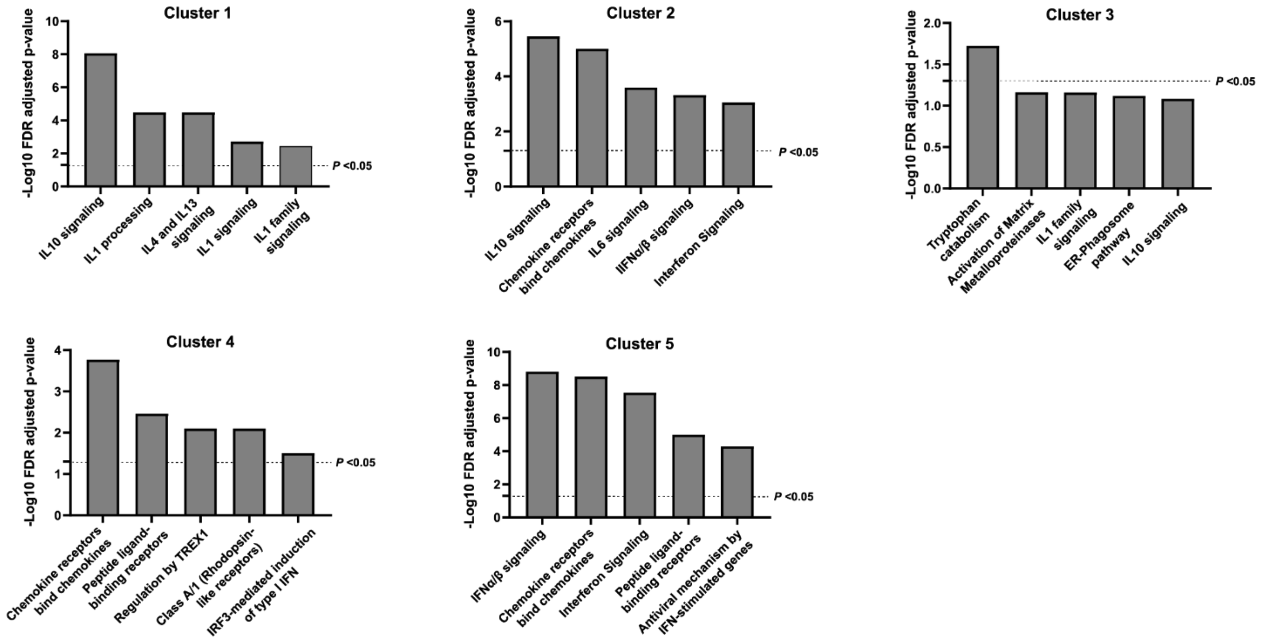


**Figure S4.**

**Reactome Pathway Analysis for donor $D^{low}$.** Histograms illustrating the five most significant pathways and the associated False Discovery Rate (FDR) corrected P-values for the gene clusters derived from the unsupervised hierarchical clustering analysis displayed in Fig 4D for $D^{low}$. Reactome uses the Binomial Test to assess the probability that the overlap between the query and the pathways has occurred by chance and the FDR is calculated using the Benjamini-Hochberg approach. Dotted lines indicate statistical significance.

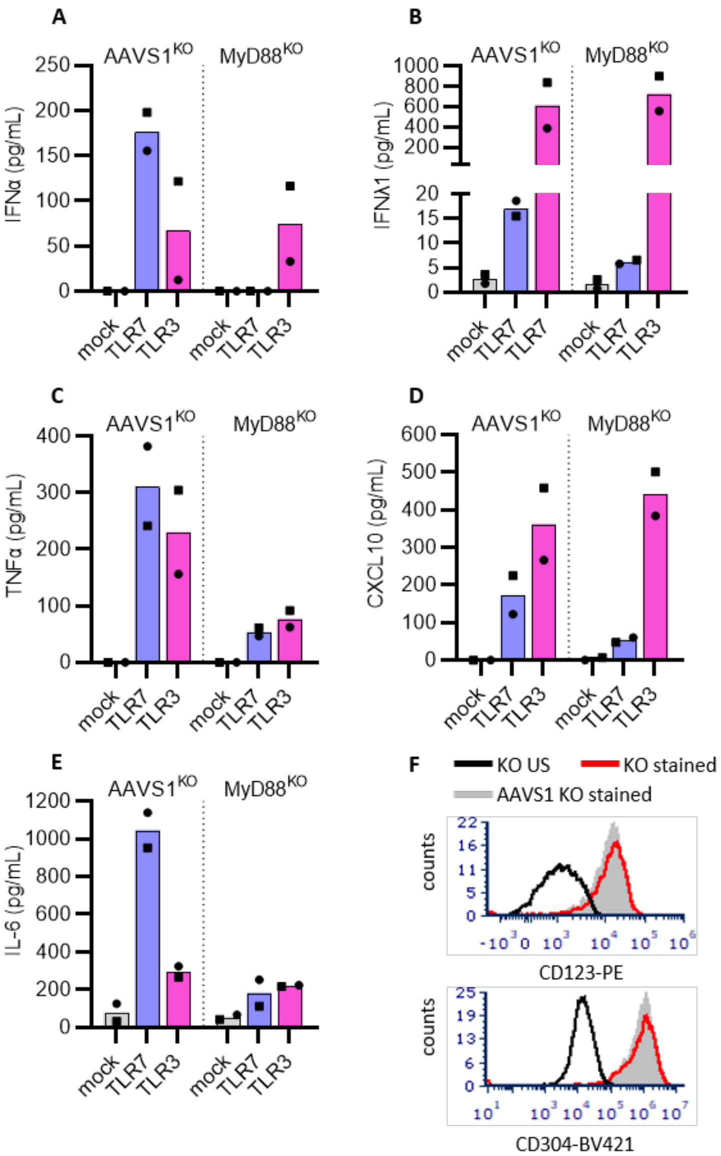


**Figure S5.**

**Validation of MyD88$^{KO}$ pDCs.** MyD88$^{KO}$ pDCs were functionally evaluated by stimulating the cells with TLR7 (2.5 µg/mL R837, blue) or TLR3 (800 ng/mL poly(I:C), pink) agonist and collecting supernatant 24 hrs after stimulation to quantify IFNα (A), IFNλ1 (B), TNFα (C), CXCL10 (D) and IL-6 (E) protein concentrations. MyD88$^{KO}$ pDC were phenotypically evaluated for expression of the pDC markers CD123 and CD304, and compared to unstained (US) MyD88$^{KO}$ pDCs and stained HSPC-pDC-AAVS1$^{KO}$ (F).

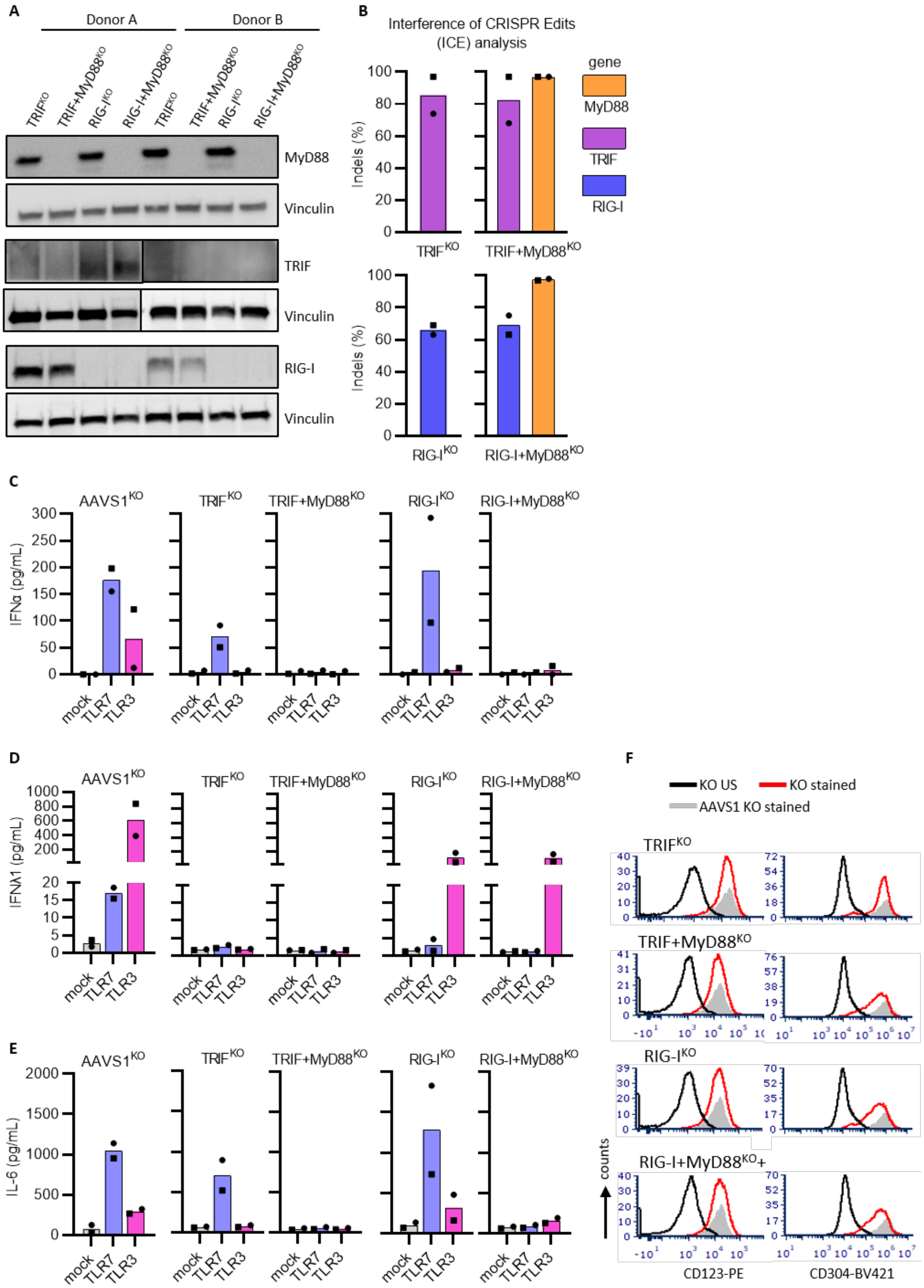
A
Donor A
Donor B
TRIF^KO
TRIF+MyD88^KO
RIG-I^KO
RIG-I+MyD88^KO
MyD88
Vinculin
TRIF
RIG-I
B
Interference of CRISPR Edits (ICE) analysis
Indels (%)
gene
MyD88
TRIF
RIG-I
C
AAVS1^KO
IFNα (pg/mL)
mock
TLR7
TLR3
D
IFNλ1 (pg/mL)
E
IL-6 (pg/mL)
F
KO US
KO stained
AAVS1 KO stained
counts
CD123-PE
CD304-BV421

**Figure S6.**

**Validation of TRIF$^{KO}$ and RIG-I$^{KO}$ pDCs.** Knock out efficiency was evaluated at the protein level by western blot analysis (A) for MyD88 (top panel), TRIF (middle panel) and RIG-I (bottom panel), and by genomic sequencing and ICE analysis to assess the frequency of insertion and deletions (indels) at the targeted sites (B). The TRIF antibody functionality was confirmed with THP-1 cell lysates (data not shown), yet we were unable to detect TRIF protein in donor B. Additionally, pDC$^{KO}$ cells were evaluated functionally by stimulating the cells with TLR7 (2.5 µg/mL R837, blue) or TLR3 (800 ng/mL poly(I:C), pink) agonist and collecting supernatant after 24 hrs to quantify IFNα (C), IFNλ1 (D) and IL-6 (E) protein concentrations. pDC$^{KO}$ were phenotypically evaluated for expression of the pDC markers CD123 and CD304, and compared to unstained (US) KO cells and stained control AAVS1$^{KO}$ pDC (F). Bars represent mean values and equal symbols represent equal donors (n=2, donor A is represented by the circle and Donor B by the square symbol).

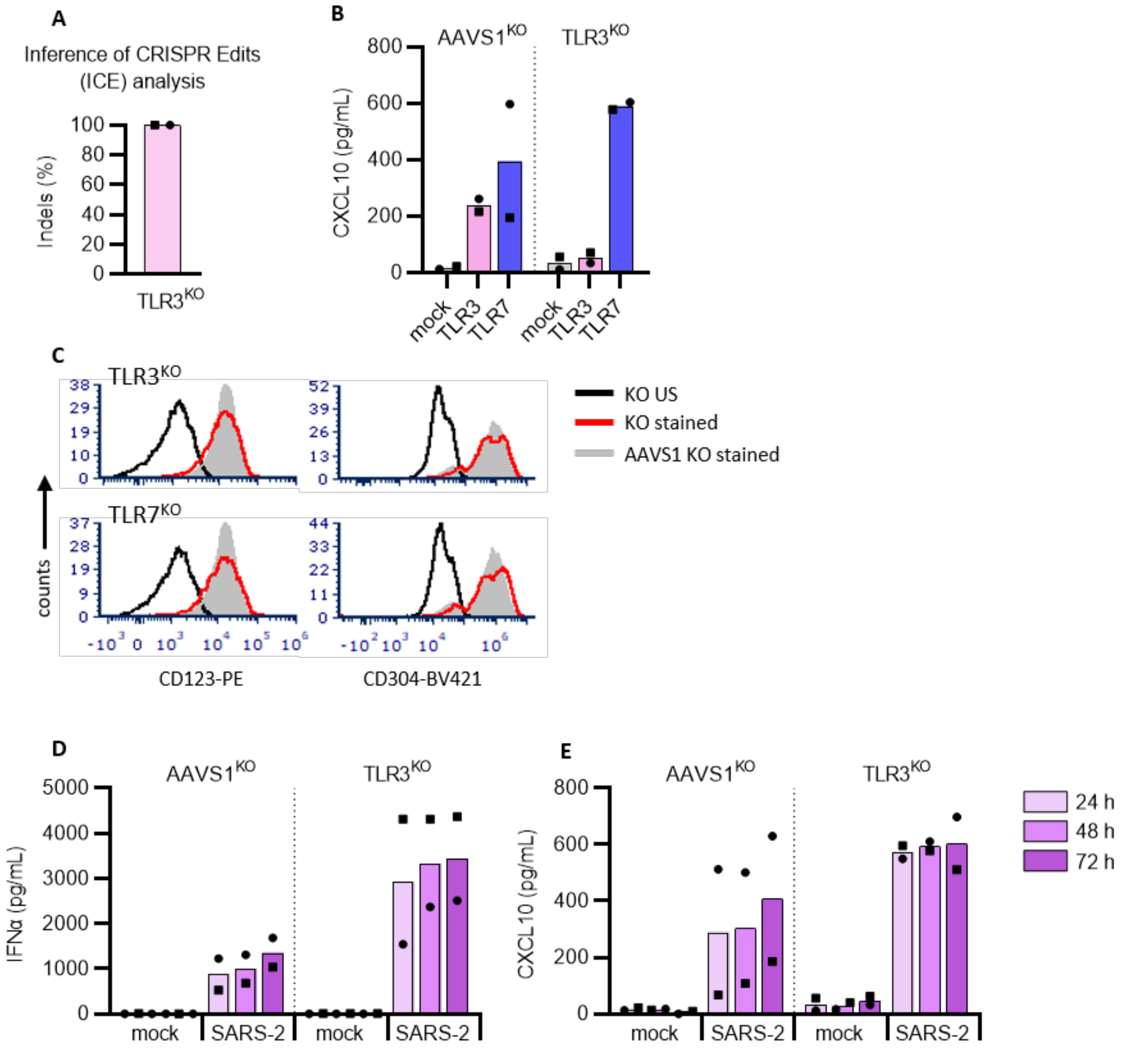


**Figure S7.**

**TLR3 is not involved in SARS-CoV-2 mediated type I IFNα production by pDCs.**

Using CRISPR/Cas9, TLR3 knock-out (KO) and AAVS1$^{KO}$ (control) pDCs were generated and cellular DNA was sequenced for ICE analysis (A). For functional evaluation, the KO pDCs were stimulated with TLR7 (2.5 µg/mL R837, blue) or TLR3 (800 ng/mL poly(I:C), pink) agonist, supernatant was collected after 24 hrs and analyzed for CXCL10 protein expression by ELISA (B). Phenotypic evaluation of pDC$^{KO}$ by flow cytometry (C). Histograms showing the expression of pDC markers CD123 and CD304, as compared to unstained KO pDCs and stained AAVS1$^{KO}$ control pDCs. AAVS1$^{KO}$ and TLR3$^{KO}$ pDCs were either mock treated (mock) or exposed to SARS-CoV-2 (SARS-2, 1 MOI), supernatants were collected at indicated time points and analyzed for type I IFNα (D) and CXCL10 (E) proteins. Bars represent mean values and equal symbols represent equal donors (n=2).

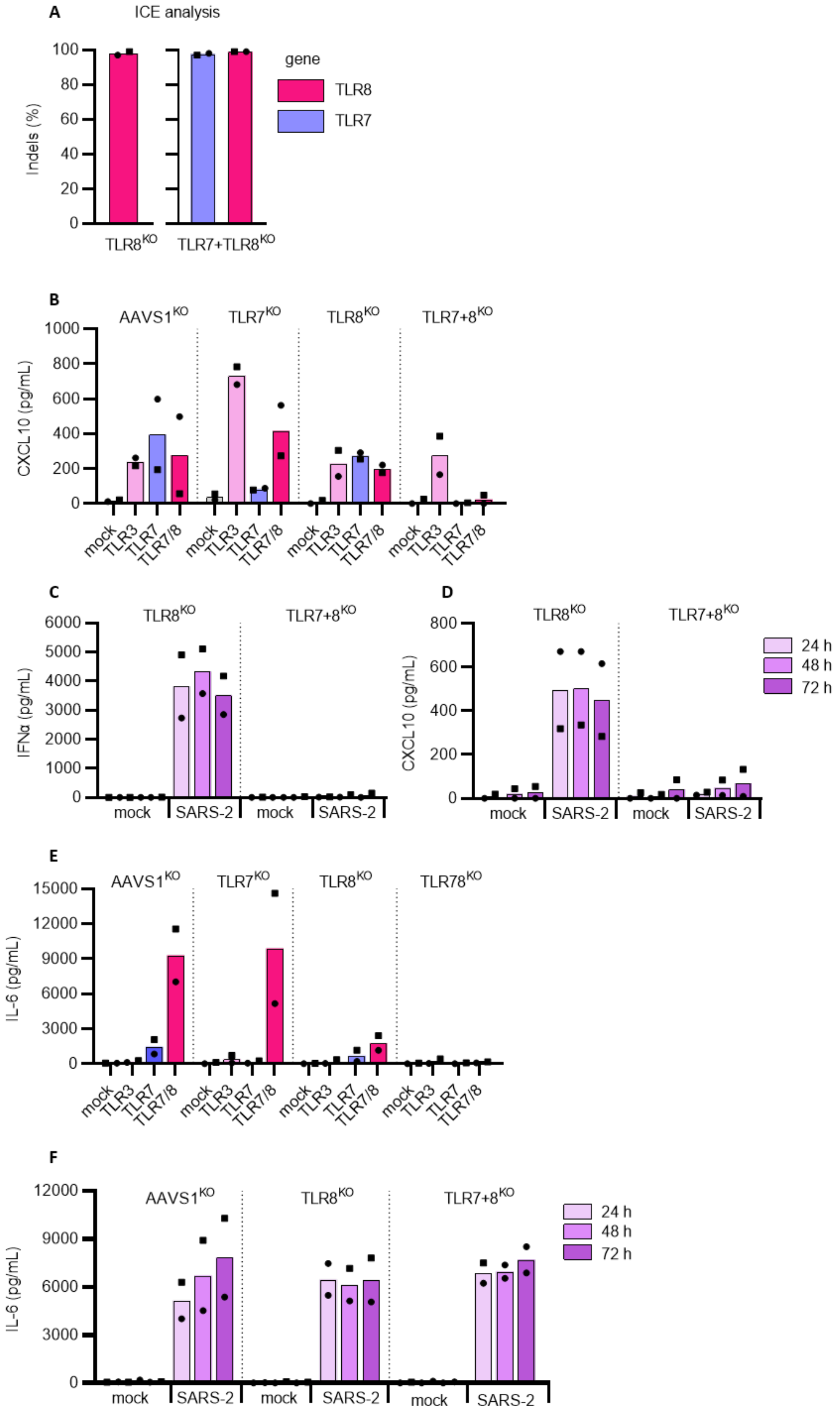
A
ICE analysis
Indels (%)
100
80
60
40
20
0
TLR8KO
TLR7+TLR8KO
gene
TLR8
TLR7
B
AAVS1KO
TLR7KO
TLR8KO
TLR7+8KO
CXCL10 (pg/mL)
1000
800
600
400
200
0
mock
TLR3
TLR7
TLR7/8
C
TLR8KO
TLR7+8KO
IFNα (pg/mL)
6000
5000
4000
3000
2000
1000
0
mock
SARS-2
D
TLR8KO
TLR7+8KO
CXCL10 (pg/mL)
800
600
400
200
0
24 h
48 h
72 h
E
AAVS1KO
TLR7KO
TLR8KO
TLR78KO
IL-6 (pg/mL)
15000
12000
9000
6000
3000
0
F
AAVS1KO
TLR8KO
TLR7+8KO
IL-6 (pg/mL)
12000
9000
6000
3000
0
mock
SARS-2

**Figure S8.**

**TLR8 is not involved in SARS-CoV-2 mediated type I IFNα production by pDCs.** To determine if the intracellular RNA sensor TLR8 can detect SARS-CoV-2 and induce cytokine production, $TLR8^{KO}$ and $TLR7+8^{KO}$ pDCs were generated and inference of CRISPR Edits was analyzed by sequencing (A). To functionally evaluate the KOs, $AAVS1^{KO}$, $TLR7^{KO}$, $TLR8^{KO}$ and $TLR7+8^{KO}$ pDCs were either left untreated (mock, grey) or stimulated with TLR3 (800 ng/mL poly(I:C), pink), TLR7 (2.5 µg/mL R837, blue) or TLR7/8 (2.5 µg/mL R848, red) agonist, supernatant was collected after 24 hrs and analyzed for CXCL10 protein expression by ELISA (B). $TLR8^{KO}$ and $TLR7+8^{KO}$ pDC were either mock treated or exposed to SARS-CoV-2 (1 MOI) and cell culture supernatants were analyzed for type I IFNα (C) and CXCL10 (D) protein production at indicated time points. To investigate TLR8-driven IL-6 production, $AAVS1^{KO}$, $TLR7^{KO}$, $TLR8^{KO}$ and $TLR7+8^{KO}$ pDCs were either treated with the different TLR agonist (E) or SARS-CoV-2 (F) and IL-6 was quantified in cell culture supernatant after 24 hrs (E) or at indicated time points (F) by ELISA. Bars represent mean values and equal symbols represent equal donors (n=2).

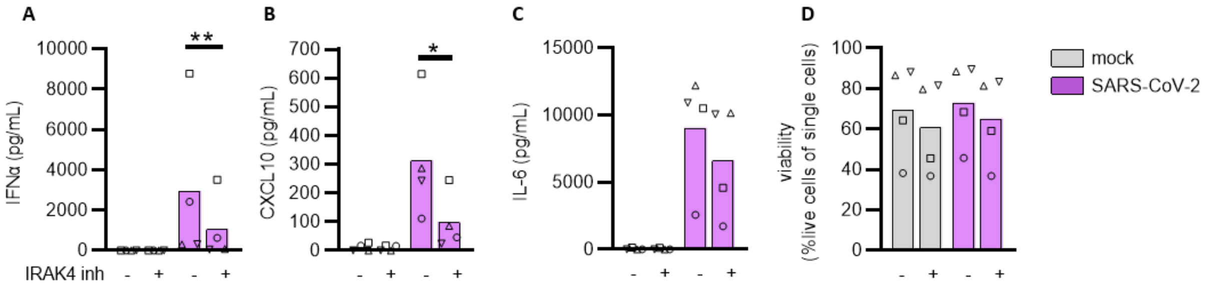


**Figure S9.**
**Inhibition of IRAK4 decreases SARS-CoV-2-induced production of type I IFNα and CXCL10 but not IL-6 by pDCs.** PDC were exposed to SARS-CoV-2 (0.5 MOI) in the absence or presence of an IRAK4 inhibitor (10 μM), and 24 hrs after virus exposure cell culture supernatants were harvest and analyzed for secretion of type I IFNα (A), CXCL10 (B) and IL-6 (C) proteins. Cells were also analyzed for viability by flow cytometry (D). Bars represent mean values and equal symbols represent equal donors (n=4). Statistical significance was determined using the ratio paired student T test. *<p0.05, **<p0.01.

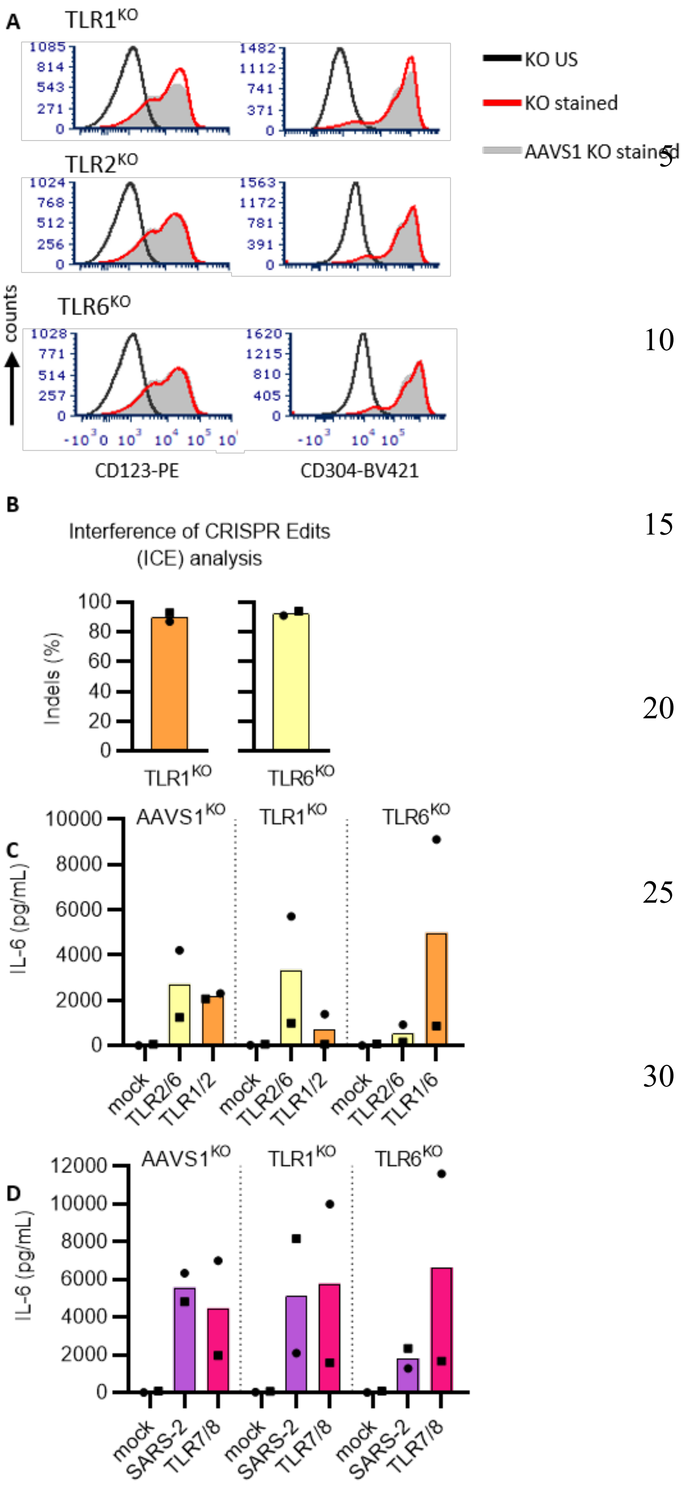


**Figure S10.**
**Validation of TLR1$^{KO}$, TLR2$^{KO}$ and TLR6$^{KO}$ pDCs.** A: Phenotypic evaluation of pDC$^{KO}$ by flow cytometry. Histograms showing the expression of pDC markers CD123 and CD304, as compared to unstained KO pDCs and stained AAVS1$^{KO}$ control pDCs. B: KO pDCs were evaluated by genomic sequencing followed by inference of CRISPR edits (ICE) analysis to assess the frequency of insertions and deletions (indels) at the target site. C: Functional evaluation the TLR1$^{KO}$ and TLR6$^{KO}$ pDCs by stimulating with TLR2/6 (5 ng/mL Pam2CSK4, yellow) or TLR1/2 (50 ng/mL Pam3CSK4, orange). Supernatant was collected after 24 hrs and analyzed for IL-6 protein expression by ELISA. D: TLR2 forms heterodimers with TLR1 and TLR6, and thus AAVS1$^{KO}$, TLR1$^{KO}$ and TLR6$^{KO}$ pDCs were either mock treated (grey), exposed to SARS-CoV-2 (0.5 MOI) or TLR7/8 agonist (2.5 µg/mL R848, red) and IL-6 protein concentrations were quantified in cell culture supernatant after 24 hrs. Bars represent mean values and equal symbols represent equal donors (n=2).

**Table S1.**

**Pathways covered by the NanoString nCounter analyses.**

Table S1 is provided as a separate excel file named: van der Sluis et al Science Report_Table S1

**Table S2.**

**List of genes that were expressed above background and subsequent analysis for $D^{high}$.**

Table S2 is provided as a separate excel file named: van der Sluis et al Science Report_Table S2

**Table S3.**

**List of genes that were expressed above background and subsequent analysis for $D^{low}$.**

Table S3 is provided as a separate excel file named: van der Sluis et al Science Report_Table S3
